# Perturb-seq identifies co-regulated gene programs shaping hematopoietic stem and progenitor cell function

**DOI:** 10.64898/2026.08.27.747033

**Authors:** Joseph Bowness, Aina Bernal Martínez, Jan Bařinka, Jonas Schulte-Schrepping, Simon Renders, Alexander Waclawiczek, Aino-Maija Leppä, Andreas Trumpp, Simon Raffel, Simon Haas, Lars Velten

## Abstract

To sustain blood formation, hematopoietic stem and progenitor cells (HSPCs) coordinate a multitude of cell biological processes, from cell cycle control and stress responses to lineage priming. While many genetic regulators of high-level HSPC function have been identified, how HSPCs coordinate more basal cell biological programs, and how such programs relate to stem cell function, remains incompletely understood. Here we use Perturb-seq to profile the transcriptional consequences of targeting 520 genes by CRISPRi in primary mouse HSPC cultures. We developed an analytical strategy to separate perturbation-induced changes in cell-state abundance and clonal heterogeneity from cell-state-local transcriptional effects. From these local perturbation signatures, we identified 19 gene regulatory programs (GRPs) that are defined by co-regulation in response to genetic perturbation, in contrast to co-expression or human curation, and align well with cell biological processes. By decomposing gene expression data from functional and clinical studies into program activity, we show that GRP activities associate with, and predict, phenotypes such as clonal output after transplantation, as well as survival and drug response in retrospective acute myeloid leukemia (AML) cohorts. Together, our study establishes perturbation-derived co-regulation programs as an interpretable framework for linking genetic regulators, cell-biological processes and stem-cell-associated phenotypes.

## Introduction

A multitude of molecular and cellular processes underpin biological function. In the context of life-long blood production, a pool of hematopoietic stem and progenitor cells (HSPCs) need to tightly control and balance the cell cycle, quiescence, symmetric vs. asymmetric division, cell growth, lineage priming, self-renewal, response to external signals, as well as niche localization and mobilization programs, among others. Changes in any of these programs can affect higher-order biological function, such as performance in transplantation assays, hematological dysfunction in advanced age, or outcome in hematological disease. In the context of HSPCs, most literature focuses on the genetic regulation of higher-order functions: this includes genome-wide association studies of blood traits^1^, classical dissection of gene functions in mouse models^2,3^, and more recent CRISPR screens of differentiation outcomes and function in transplants^4–6^. The molecular regulation of elemental cellular processes in HSPCs, and how these processes collectively mediate higher-order functions, remains less well described, and is characteristic of a broader gap in bridging gene regulatory programs to complex biological phenotypes^7^.

Functional genomics approaches enable systematic dissection of the molecular regulation of cellular processes. In particular, the combination of CRISPR screens and single-cell RNA-seq (“Perturb-seq”, also known as CROP-seq, CRISP-seq etc.) facilitates characterization of transcriptomic programs and their regulators by combining specific genetic perturbations with a genome-wide transcriptional readout^8–11^. Perturb-seq can identify both groups of genes that elicit similar effects when perturbed (“co-regulating” genes: often genes that interact closely, for example in the same protein complex or signaling pathway) and genes that respond coherently to various perturbations (“co-regulated” genes)^12–16^. Perturb-seq data therefore has the potential to distinguish between genes that are co-expressed (e.g. along differentiation trajectories) and genes that exhibit bona fide co-regulation across different cell states and contexts. While co-expression programs are widely studied, few studies have exploited co-regulation signatures^17^. In the hematology field, gene expression signatures have a substantial value for interpreting transcriptome data^4,18,19^ and predicting clinical outcomes in diseases such as acute myeloid leukemia^20–22^, but the value of co-regulation programs for these tasks has not been investigated.

Previous Perturb-seq studies^8–10,12,13,16^ have mostly been conducted in cell lines and large-scale datasets from primary cells are scarce. In hematopoiesis, Perturb-seq has largely been constrained to characterizing differentiation outcomes (e.g. upon transplantation) rather than profiling molecular programs of HSPCs^5,11,23^. Recent advances in culture protocols^24,25^ that facilitate the maintenance and expansion of functional hematopoietic stem cells (HSCs)^26,27^ now provide an opportunity to perform Perturb-seq screens in large numbers of HSPCs isolated from animals. In this study, we performed a CRISPR interference (CRISPRi) Perturb-seq screen for over 500 chromatin-associated regulators of hematopoiesis in primary mouse HSC expansion cultures^24^. We developed analysis routines to separate biological noise, perturbation-induced differentiation state changes, and local consequences of perturbations by comparing perturbed cells to unperturbed controls within cell states. The latter grouped perturbations into regulatory complexes and served to identify genes coherently co-regulated across perturbations. Of these 19 co-regulation signatures, which we term gene regulatory programs (GRPs), at least 13 were strongly associated with well-defined cell biological processes, ranging from translation initiation to signaling pathways and lineage priming. To associate elemental GRPs with higher-order stem cell function, we decomposed RNA-seq data from human and mouse studies of HSPC and leukemic stem cell function into GRP activity. We show that GRP activities capture and predict functional variation in hematopoietic stem cell self-renewal and engraftment, and improve prediction of long-term survival and Venetoclax response in acute myeloid leukemia (AML) beyond other transcriptome-based measures including co-expression programs. In sum, our work introduces programs of genetic co-regulation as an interpretable framework for linking genetic regulation, HSPC cell biology and stem-cell-associated phenotypes.

## Results

### Perturb-seq in cultures of primary hematopoietic stem and progenitor cells (HSPCs)

In order to effectively infer genetic programs of co-regulation, we needed to measure transcriptional variation in HSPCs caused by a broad range of perturbations to key regulators. To generate this data, we used CRISPRi coupled with single-cell RNA sequencing of perturbed transcriptomes (Perturb-seq). We selected a total of 520 genes to target for knockdown by a ranking system which integrated expression metrics and prior literature implicating genes in hematopoiesis, with particular focus given to transcription factors and other chromatin-associated regulators (Figure 1A, Table S1, Methods). Three single guide RNAs (sgRNAs; hereafter, ‘guides’ or ‘guide RNAs’) per target gene were cloned as a pool into a lentiviral CROP-seq-GFP vector^10,28^ (Figure S1A). We also included 120 negative control guides - comprising approximately 10% of the total library - of either non-targeting spacer sequences or sequences targeting a single intergenic location in the genome (Figure 1A).

**Figure 1.**
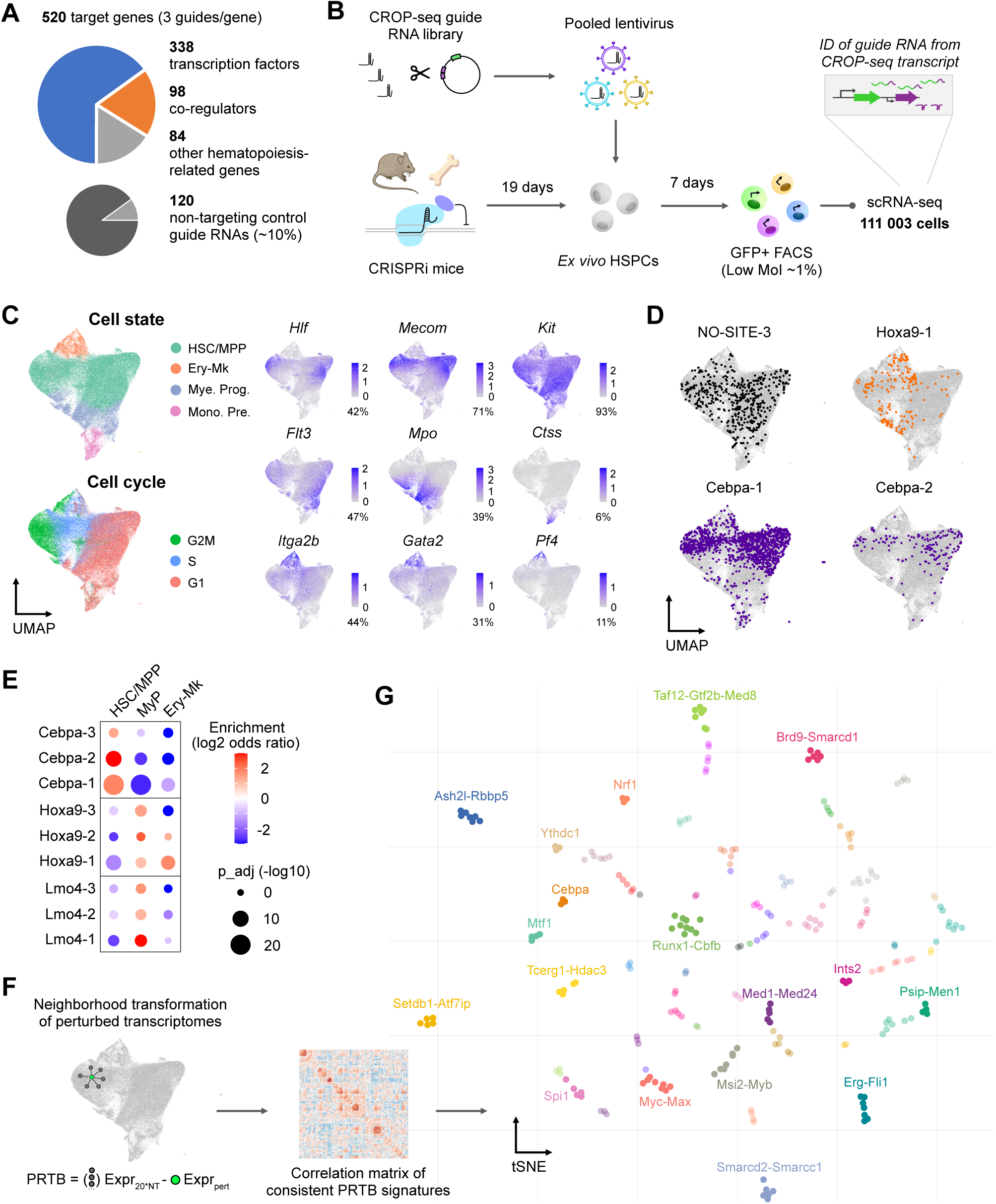
Perturb-seq of 520 target genes in HSC expansion cultures. **A.** Pie chart categorizing genes targeted by CRISPRi. 120 non-targeting control guides represent ∼10% of the total guide RNA library. **B.** Experimental design of Perturb-seq in HSC expansion cultures. See Figure S1 for flow cytometry characterization of the culture system and Methods for more detailed protocols. **C.** UMAP embeddings of single cells from the Perturb-seq experiment, annotated by cell state and cell cycle labels. Cell state was annotated from marker gene expression, with some examples shown (Hematopoietic stem cells and multipotent progenitors: *Hlf*, *Mecom, Kit, Flt3;* Myeloid progenitors: *Mpo*; Monocyte precursors: *Ctss;* Erythroid progenitors: *Gata2, Itga2b;* Megakaryocyte progenitors*: Pf4*). **D.** Examples of distribution patterns of cells containing guide RNAs over the cell state landscape. **E.** Differential abundance results for three perturbations which show enrichment/depletion over cell states in the data. **F.** Left panel: Schematic of the PRTB transformation, which subtracts the transcriptome of each perturbed cell from its nearest neighbors in cell state space. Right panel: Pairwise Spearman correlation matrix calculated from “pseudobulk” perturbation signatures of neighborhood-transformed transcriptomes. **G.** tSNE embedding of perturbations by similarity of effects on the HSPC transcriptome (co-regulat*ing* genes). Points are colored according to HDBSCAN clusters and labelled by the dominant target genes per cluster (see Table S2 for full annotation of guides in each cluster).

An overview of the Perturb-seq experimental design is presented in Figure 1B. Lineage-cKit+ cells were extracted from limb bone marrow of transgenic mice engineered for hematopoietic-specific expression of a CRISPRi repressor (LSL dCas9-KRAB^29^ × Vav1-Cre^30^) and expanded *ex vivo* under conditions optimized for HSC maintenance and expansion^24^. Flow cytometry characterization of these cultures consistently shows a Lineage-cKit+ Sca1+ HSPCs fraction of ∼60% (Figure S1B,C), as well as expression of the HSC surface marker EPCR/CD201 in >20% of cells in the culture (Figure S1C). After a 19-day expansion period, HSPCs were infected with the lentivirus-packaged CRISPRi guide RNA library at a low multiplicity of infection (MOI). Cells were cultured for a further seven days before GFP-positive cells harboring the integrated CROP-seq cassette were isolated by fluorescence-activated cell sorting (FACS) (Figure S1D). We performed single-cell RNA sequencing by split-pool combinatorial barcoding (Parse Evercode WT kit) and recovered transcriptomes from a total of 111,003 cells after quality control filters.

We assigned guide RNAs to cells by targeted amplification of CROP-seq transcripts with a comparable efficiency to other Perturb-seq studies in primary cells using alternative scRNA-seq platforms^23,31^: 79,192 of cells were assigned a single guide (71.3%), 23,644 no guide (21.3%), and 8,167 multiple guides (7.4%) (Figure S1E). This recovered an average of 188.1 cells per guide RNA (quartiles: 69.25 - 229.5) for 20 priority gene targets included at five-fold representation, and 40.65 cells per guide RNA (quartiles: 9.0 - 56.0) for guides targeting the remaining genes. CRISPRi was broadly effective (386/520 targets significantly repressed, mean log2 fold change = -1.35) with repression variable by target gene but relatively consistent between the three guide RNAs per gene (Figure S1F). We further performed an essentiality analysis by comparing guide distributions in the CROP-seq vector with cell numbers recovered in the Perturb-seq data set. This analysis identified a suite of known essential genes whose knockdown impairs survival/proliferation (e.g. *Smc1a*, *Top2a*, *Ruvbl2*) (Figure S1G). Together, these analyses demonstrate the successful implementation of Perturb-seq in HSC expansion cultures.

### Identifying perturbation effects in heterogenous HSPC cultures

We next characterized the cell state landscape in the culture conditions from the transcriptomic profiling of the Perturb-seq data (Figure 1C). Whilst ∼40% of cells demonstrate expression of canonical HSC marker genes such as *Hlf* and *Mecom*, the data also contains a substantial fraction of early myeloid-biased progenitors (*Mpo*+), as well as smaller subpopulations of erythroid-megakaryocyte progenitors and monocyte precursors. The fraction of transcriptome-defined HSCs we observed is similar to that reported in other studies also using 5% O_2_, while exceeding that achieved by studies adapting the original Wilkinson et al expansion protocol at 20% O_2_ (Figure S2). Cell cycle differences constitute an additional axis of intercellular heterogeneity within the data (Figure 1C).

One way perturbations effects manifest is through altered cellular differentiation, leading to enrichment or depletion of perturbed cells in certain cell states compared to controls (Figure 1D). We identified perturbations causing differential abundance in the cell states represented within our data that were consistent over multiple guide RNAs (Figure S3A). For example, *Cebpa*-targeted cells were consistently enriched in HSPCs and depleted from early myeloid-biased progenitors, while *Hoxa9* and *Lmo4* knockdown cells were depleted from HSPCs and more abundant in myeloid progenitors (Figure 1E).

Besides the biological effects of perturbations on cell state, we noticed some significant biases unrelated to genetic perturbation, for example for two of the 120 non-targeting control guides (Figure S3B). These patterns likely arise from clonal proliferation of limited numbers of progenitor cells during the 7 days in culture between lentiviral infection and transcriptome profiling, occurring despite the anticipated infection of hundreds of cells per guide in our experimental design. This confounding challenge of clonal heterogeneity in Perturb-seq has also been reported in other contexts such as embryoid body differentiation^32^.

To produce a resource of differential gene expression results for all perturbations in our experiment (see *Data availability*), we accounted for clonal heterogeneity by calibrating our analysis using the SCEPTRE package^33^ and quantifying consistency across the three guides targeting each gene (Supplementary Note, Figure S3C,D). A smaller biological replicate of the screen demonstrates these results are reproducible, at least for perturbations covered by sufficient cell numbers in the replicate screen (Supplementary Note, Figure S3E-G). Notably, differential expression analysis in this experimental context of 7 days of CRISPRi encompasses both the direct target genes of perturbations and secondary transcriptional consequences of effects on cell state.

### Neighborhood transformation masks heterogeneity to reveal co-regulating genes in HSPCs

We next aimed to focus on local, within-state gene expression changes induced by perturbations, separating them out from cell state-related effects. As a strategy to achieve this separation, we implemented the ‘perturbation signature’ (PRTB) transformation^34^, which calculates the difference of each perturbed cell’s transcriptome from the average of its 20 nearest neighbor control cells (Figure 1F). These neighborhood-transformed gene expression vectors thus represent perturbation-specific differences compared to unperturbed cells of a similar cell state. Practically, applying the PRTB transformation to our Perturb-seq data suppressed cellular variability related to differentiation state and cell cycle (Figure S4A), along with biased patterns of guides over the cell state landscape (Figure S4B).

When we aggregated PRTB scores across all cells containing each guide, the neighborhood transformation reduced noise in the matrix of pairwise correlations to more clearly reveal signatures of perturbations correlated between the three guides targeting the same gene (Figure S4C). We then also aggregated PRTB scores separately in different cell states, and found that perturbation signatures were highly correlated whether calculated using cells from all states, or only HSCs/MPPs or Hlf+ cells (Figure S4D). By contrast, PRTB scores calculated from more differentiated progenitor populations were less correlated. These results suggest that the signatures computed here mostly capture regulatory relationships at a stem cell and early progenitor level.

After filtering the data set to perturbations where effects were consistent between guides targeting the same gene (see Methods: *Filtering for consistent perturbations* and Figure S4E) (n = 246/1560 guide perturbations), we could embed perturbations by the similarity of their transcriptome-wide effect, thus associating co-regulati*ng* genes (Figure 1G). In this embedding, guides targeting the same gene (Figure S5) and, more notably, genes with well-known biochemical interactions clustered together (Figure 1G). For example, Runx1 and Cbfb form a heterodimer to bind DNA^35^, Max is a co-factor of Myc^36^, and Psip1/Ledgf is a binding partner of Menin^37^. We also observed clusters representing chromatin-associated complexes such as SET1/MLL (Ash2l-Rbbp5), canonical BAF (Smarcc1-Smarcd2), and non-canonical BAF (Smarcd1-Brd9). Interestingly, the transcription factor Myb clustered with the RNA-binding protein Msi2. Myb RNA has previously been identified in the Msi2 interactome^38^ and Msi2 levels have been identified as a genetic regulator of HSC self-renewal^39^ and clonal hematopoiesis risk^40^. Our data thus implies that Msi2 regulation in HSPCs is mediated through interplay with Myb.

Altogether, these analyses demonstrate that local gene expression signatures of perturbations group perturbations by protein complex, suggesting that they capture programs of co-regulation in HSPCs.

### Co-regulated gene sets define elemental gene regulatory programs (GRPs)

Next, we used our Perturb-seq data set to link perturbations to cellular processes by the inference of co-regulat*ed* gene programs. Specifically, we applied Guided Sparse Factor Analysis (GSFA)^41^ on neighborhood-transformed perturbation signatures to identify weighted gene sets (factors) that coherently change across perturbations. By combining (a) Perturb-seq-induced variation, (b) suppression of unrelated cell-state variability by the PRTB transformation, and (c) perturbation-guided factor inference (GSFA), this framework is particularly well suited to detect co-regulation, in contrast to other single-cell RNA-seq factor analysis methods such as Non-negative Matrix Factorization (NMF), which primarily capture sets of genes co-expressed across cell states. The use of neighborhood-transformed perturbation signatures for downstream factor inference also reduces the impact of clone-associated guide biases compared to differential expression analysis of individual perturbations.

Based on statistical considerations (Figure S6A), we ran GSFA with K=21 components, although inferred factors were consistent across different K values (Figure S6B). This resulted in 19 non-empty factors in the form of relatively small lists of mostly non-overlapping genes (median: 89 genes, Figure 2A, Table 1), hereafter termed gene regulatory programs (GRPs). Of these GRPs, at least 13 were strongly enriched for well-defined biological processes, as evidenced by results of GO term enrichment (Table 1, Figure S6C), manual examination of the genes loaded to each GRP (Table 1, Table S3), and large language model-based annotation^42^ (Table 1). In later sections, we provide further evidence that informed the functional annotation of the GRPs (see Figure 3 and 4, Table 1).

**Figure 2.**
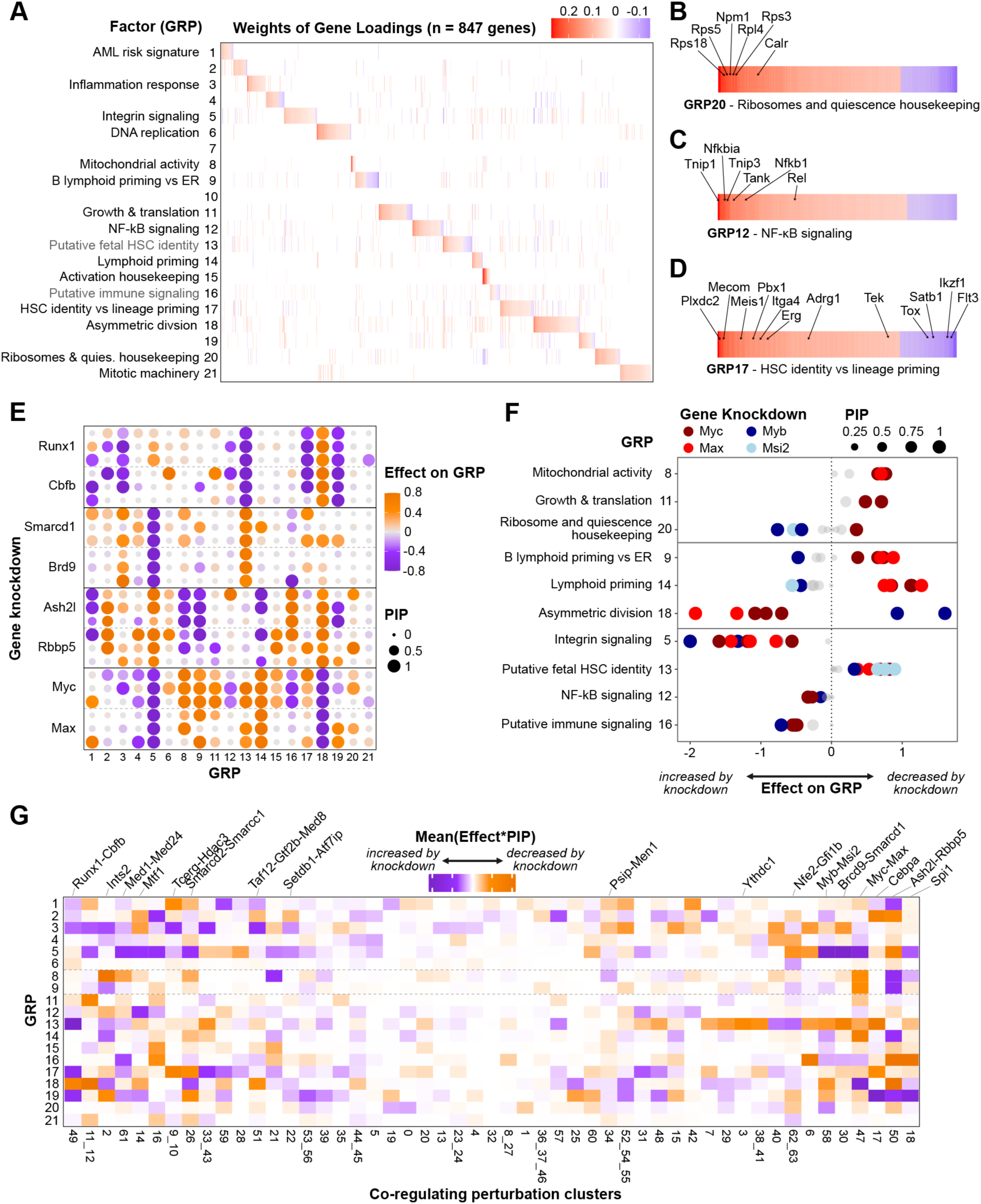
Perturbation-guided inference of elemental gene regulatory programs (GRPs) **A.** Heatmap of all genes significantly loaded onto GRPs derived by guided sparse factor analysis (GSFA), ranked and colored according to weight of gene loading. See Methods for details of GSFA implementation and Table 1 for justification of GRP names. Factors 7 and 10 had no genes loaded with an inclusion probability > 0.8 (i.e. “empty” factors), whereas factors 2, 4 and 19 could not be readily annotated with a biological process. **B-D.** Insets of individual GRPs labeling representative genes (see Table S3 for full lists of genes loaded to each GRP). **E.** Dot plot of the effects of selected perturbations (3 guides per gene knockdown) on GRPs, as inferred by GSFA. Colors of dots represent strength of effect (capped at 0.8) and sizes represent the posterior probability that the perturbation has an effect on the GRP. **F.** Dot plot illustrating effects of Myc-Max and Myb:Msi2 knockdown on selected GRPs. **G.** Heatmap of aggregated perturbation effect on GRPs for all clusters of co-regulating perturbations defined in Figure 1F. For the constituent perturbations within co-regulation clusters that are not labelled above the plot, see Table S2.

**Figure 3.**
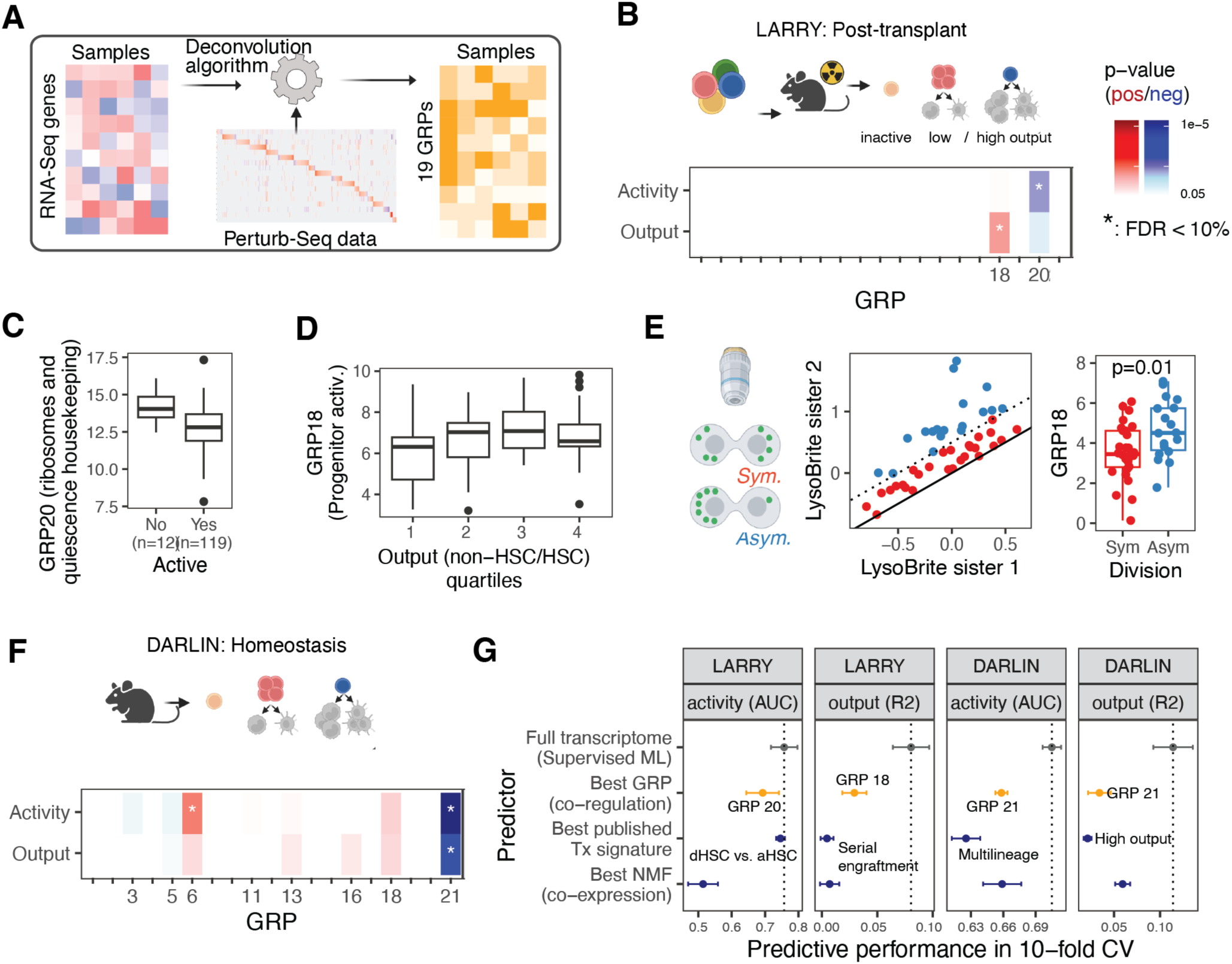
Decomposition of HSC transcriptomes links GRPs to stem cell function. For methodological detail, see Methods, *Decomposition of HSC transcriptome data into GRPs* and *Evaluation of the predictive value of GRPs* **A.** Illustration of the approach. **B.** Association of GRPs with clonal function post-transplant (LARRY)^48^. HSC transcriptomes were aggregated per LARRY clone, decomposed into GRP and the statistical association with activity (defined as presence of any non-HSCs in clone) and output (defined as ratio of non-HSCs and HSCs per clone) was computed. P-values are from a Wilcoxon test (for activity) or a correlation test (for output). **C.** Box plot illustrating activity of GRP20 in HSCs from n=131 LARRY clones, stratified by activity. **D.** Box plot illustrating activity of GRP18 in the same clones, stratified by output. **E.** Relationship of GRP18 activity to division mode. HSC transcriptomes were obtained from a data set where single-cell RNA-seq had been performed subsequent to live cell imaging and tracing of lysosome content during cell division^50^ (illustrated in the left panel). Sister cell pairs were classified as symmetrical or asymmetrical division based on lysosome distribution patterns (central panel). GRP18 activity was quantified by decomposition of single-cell RNA-seq data. Data shown is for the stem cell daughter (i.e. the daughter receiving more lysosomes^50,66^). **F.** Association of GRPs with clonal function in homeostasis (DARLIN)^51^. Like B, except that data from the DARLIN model was used. For color scale, see B. **G.** Predictive performance of different predictors on activity and output. See Methods for detail. Note that the Best GRP, best NMF and best published Tx signature were picked during cross validation to avoid “multiple hypothesis testing” issues.

**Figure 4.**
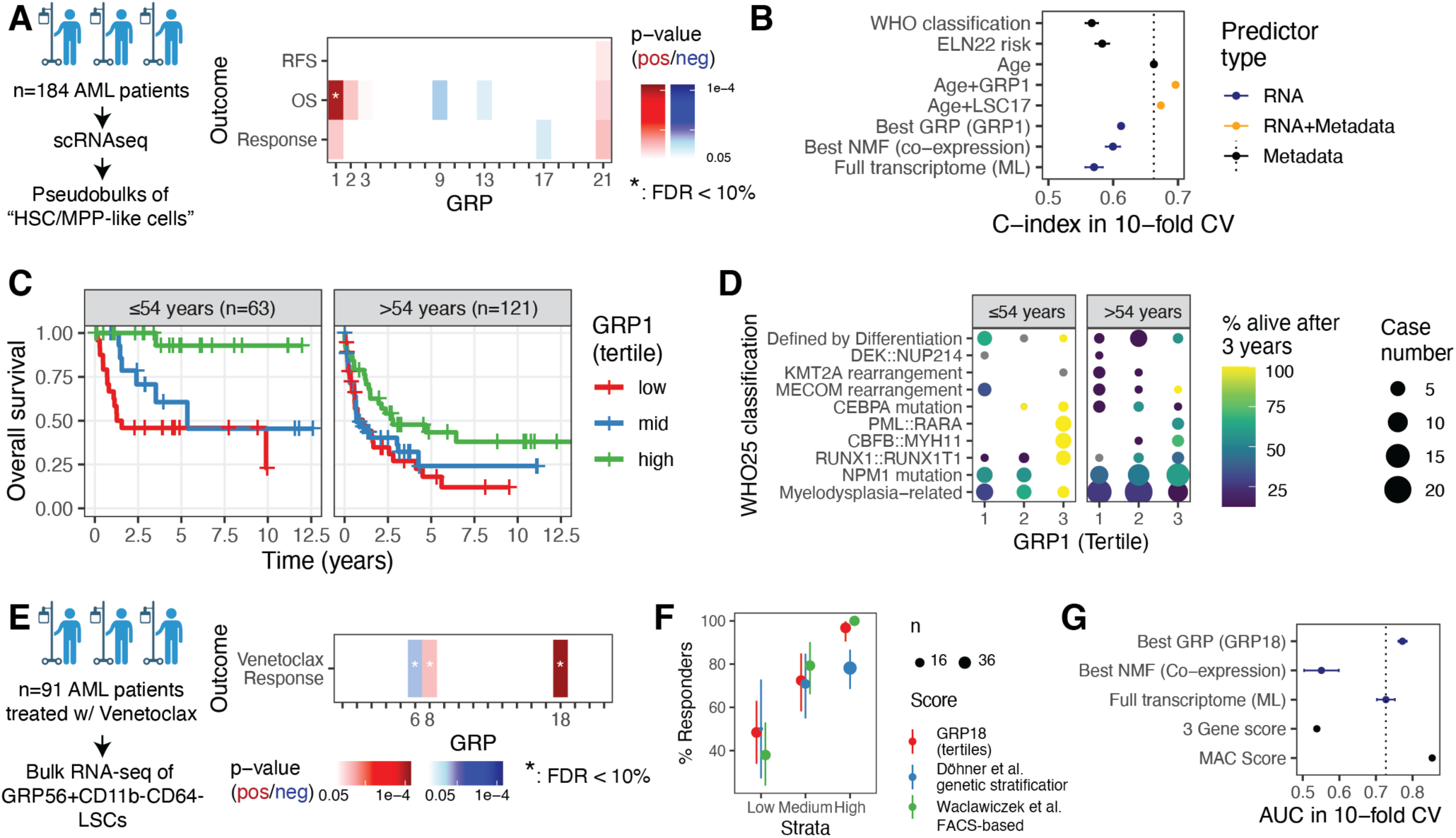
GRPs predict outcomes in AML. For methodological detail, see Methods, Decomposition of HSC transcriptome data into GRPs and Evaluation of the predictive value of GRPs. **A.** Putative LSC transcriptomes from a single-cell RNA-seq cohort were decomposed into GRPs and associations with first-line therapy response, overall survival (OS) and relapse free survival (RFS) were computed. P-values are from a Wilcoxon test (for response) or from a Cox proportional hazards regression model (for OS and RFS). **B.** Predictive performance of different predictors of OS. See Methods, section *Evaluation of predictive value of GRPs* for detail. Note that the best GRP and best NMF were picked during cross validation to avoid “multiple hypothesis testing” issues. **C.** Survival curves, stratified by age and tertile of GRP1 activity. **D.** Dot plot illustrating 3-year survival and case numbers, stratified by age and WHO classification. Cases that dropped out from follow-up in the first 3 years were excluded from the 3-year survival calculation. **E.** Putative LSC transcriptomes from a study on Venetoclax were decomposed into GRPs and association with Venetoclax response was determined. P-values are from a Wilcoxon test. **F.** Plot illustrating the percentage of responders as a function of GRP18 activity (red) or an existing genetic classification of Venetoclax response^64^ (blue). Error bars illustrate 90% confidence interval. **G.** Predictive performance of different predictors on Venetoclax response.

**Table 1.**
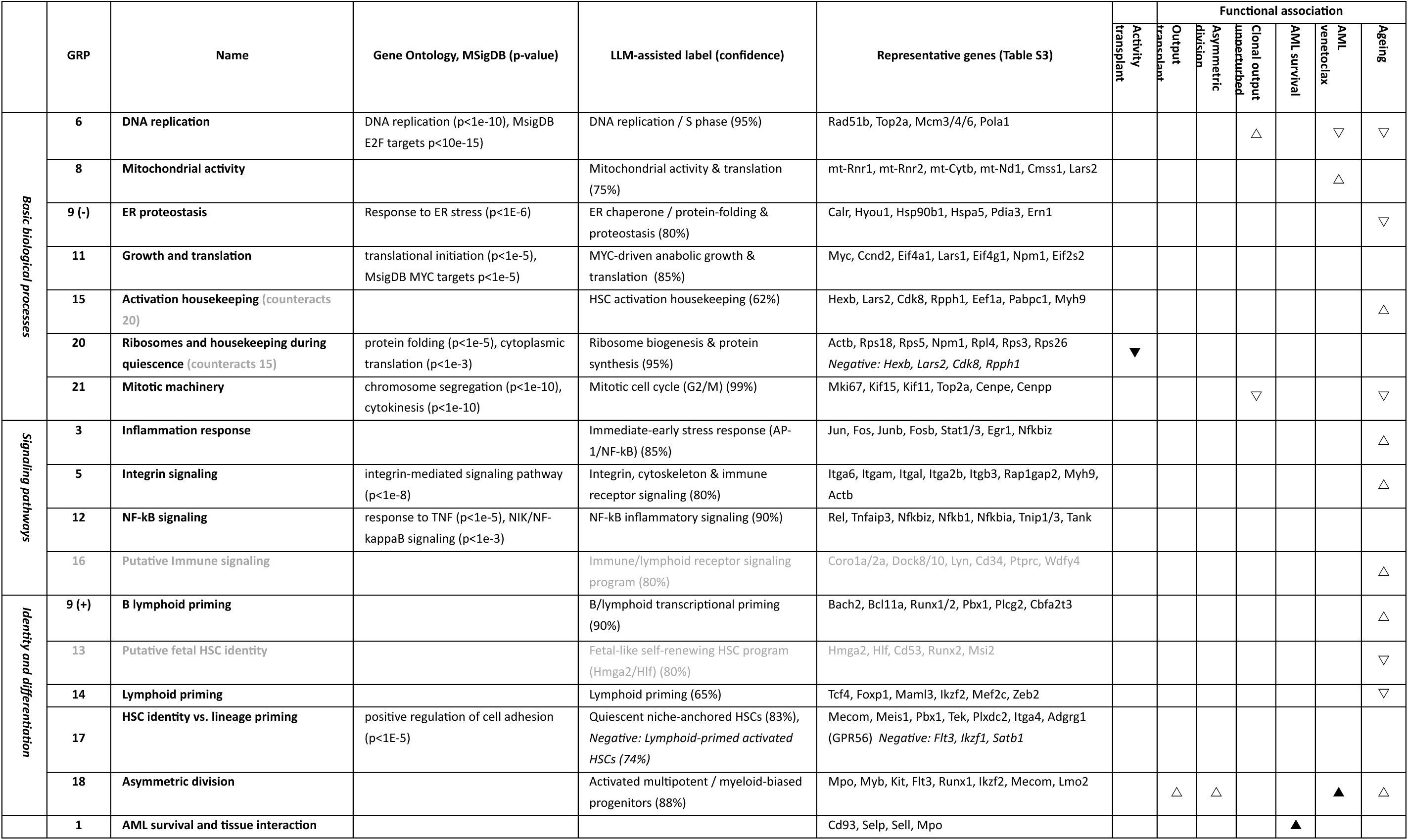
Annotation of gene regulatory programs identified from GSFA. GPT-5.1: Annotation by a large language model, see methods. Functional association: See Figure 3&4. ▾▴: strong predictive value of GRP up/downregulation on function/outcome, ▽△: statistically significant association between GRP and function/outcome

In brief, we identified GRPs related to basic cell biological processes (GRP15, GRP20: Ribosome and housekeeping programs; GRP9: *ER Proteostasis*; GRP11: *Growth and translation*; GRP6: *DNA replication*; GRP21: *Mitotic machinery*; Figure 2B) and signaling pathways (GRP3: *Inflammation response*, GRP5: *Integrin signaling*, GRP12: *NF-kB signaling*, GRP16: *Putative immune signaling*; Figure 2C). We also identified myeloid, lymphoid and B cell priming programs, and two programs associated with “stemness” (Table 1). Notably, GRPs associate each gene with a weight (loading) that can be positive or negative. In the example of GRP17, canonical HSC genes were positively loaded, whereas lymphomyeloid genes had negative loadings (Figure 2D), suggesting antagonism between these two programs across perturbations. In most cases, however, the loadings of genes had consistent signs within a GRP.

Given the inherent differences between our GRP inference methodology and factor analysis methods focused on co-expression, NMF performed on gene expression variation across unperturbed cells from our Perturb-seq data derived clearly distinct gene programs (Figure S6D). Separately, we found our approach to be largely robust to the cellular heterogeneity within the data, with most GRPs being consistently identified when we used different immature cell states as input for GSFA (Figure S6E). This result highlights the difference between programs of co-regulation and co-expression and reinforces the notion that our GRPs are mostly relevant to stem cells and early progenitors.

In sum, we identified 19 sparse programs of genes that respond consistently to genetic perturbations.

### Molecular regulators of GRPs

By its design, GSFA quantifies the effects of all perturbations on each GRP. These effects were generally consistent between replicate guide RNAs and for genes in the same regulatory clusters (Figure 2E). Some perturbations mediated effects through only a few GRPs, whilst other such as Ash2l-Rbbp5 (SET/MLL1) complexes had consistent effects on a larger number of GRPs (Figure S7A,B). As an illustrative example (Figure 2F), Myc-Max knockdown suppressed several programs in keeping with its role as a master activator of cell growth pathways, such as *mitochondrial activity* (GRP8), *growth and translation* (GRP11) and *ribosomes and quiescence housekeeping* (GRP20), and also decreases putative lymphoid priming programs (GRP9 and GRP14). By contrast, Myc-Max knockdown strongly increased *asymmetric division* (GRP18). Interestingly, Myb:Msi2 perturbation displayed opposite effects to Myc-Max on programs related to ribosome synthesis and lineage priming, but concordant effects promoting Integrin and NF-kB signaling programs (GRP5 and GRP16). These results underline that Myc and Myb regulate, in a partly opposing manner, key programs for stem cell activation^43,44^, and implicate the RNA binder Msi2 as a key mediator of the interplay between these factors. This adds to prior work reporting an important post-transcriptional component in the regulation of Myc in HSCs^45^.

The two SWI/SNF complexes represented in our data, cBAF (Smarcc1-Smarcd2) and ncBAF (Smarcd1-Brd9), also displayed some shared and some opposing effects on GRPs (Figure S7C), similar to a previous study which used Perturb-seq to interrogate a larger number of SWI/SNF components in an AML cell line^46^. More specifically, both knockdowns promoted *Integrin signaling* (GRP5) and decreased programs related to HSC identity (GRP13 and GRP17). However, only ncBAF suppressed the *inflammation response* (GRP3), whereas cBAF knockdown promoted *asymmetric division* (GRP18).

Overall, we were able to summarize perturbations of all co-regulating complexes (see Figure 1G) in terms of their effects on GRPs (Figure 2G). Thus, we provide a rich resource linking HSPC GRPs to their regulators, and demonstrate how GRPs provide a framework for interpreting how genetic perturbations mediate their effects on cells through a range of biological processes.

### Decomposition of transcriptomes into GRPs

We hypothesized that the GRPs constitute elemental units of gene regulation in HSCs and early progenitors, and therefore may be instructive of higher-order stem cell and tissue function. To investigate their relevance outside of the *ex vivo* Perturb-Seq setting, we developed a decomposition package which can be applied to any HSC/MPP transcriptome data. Our algorithm estimates ‘activity’ scores across the 19 GRPs (Figure 3A) by fitting a linear model to describe the gene expression vector of a sample as a linear combination of the factors derived from Perturb-seq. Put simply, we infer activity of the GRPs, given RNA-seq data from HSCs/MPPs.

In the following sections we apply this decomposition approach to a variety of external data sets where gene expression profiles of immature hematopoietic progenitors (HSCs or MPPs) are linked to a) clonal properties of HSCs in mouse models and b) ageing and myeloid neoplasm in humans. Of note, we caution against applying GRP decomposition to data sets that are less related to the cellular context in which the Perturb-seq was performed, i.e. the method requires HSC/MPP transcriptomes. Our algorithm is available as an R package (https://github.com/veltenlab/hsc19).

### A GRP linked to clonal stem cell output and asymmetric division

We first focused on HSC self-renewal post-transplant as an example for a biologically important functional property of HSCs. Specifically, we quantified GRP activity in a data set where clonal HSC transcriptomes were linked to clonal HSC function post-transplant (“LARRY”, Figure 3B). Here, HSCs were labeled with a lentivirus, transplanted, and both HSC transcriptomes and HSC output were determined 3 months later^4,47,48^. For each clone, we determined (i) whether the clone made any non-HSCs (“activity” as opposed to dormancy), and (ii) the “output”, defined in the original LARRY study^4^ as the ratio between non-HSCs and HSCs per clone.

We applied decomposition to the “pseudobulk” gene expression profiles of all HSCs per clone, thereby quantifying activity of all 19 GRPs per clone. We found that activity post-transplant was negatively associated with GRP20 (*Ribosomes and quiescence housekeeping,* Figure 3B,C). Interestingly, this ribosome-centric signature strongly overlapped with a previously published long-term dormant HSC signature (p<10^-16^, Figure S7D)^49^. By contrast, output post-transplant was positively associated with GRP18. This result suggests that post-transplant, some HSCs persisted in deep quiescence characterized by low expression of all programs *except* core biosynthesis machinery, whereas others entered a phase of clonal growth. Subsequently, a specific genetic program (GRP18) was associated with the balance between clonal self-renewal of HSCs and production of progenitors (Figure 3B,D). GRP18 included genes such as *Mecom*, *Flt3*, *Kit*, *Runx1*, *Myb*, and *Msi2* (Table 1, Table S3). Myc-Max and Myb:Msi2 were among the strongest regulators of that GRP (see Figures 2F, S9G).

Clonal stem cell output might be related to the balance between symmetric versus asymmetric decisions. To follow up on this hypothesis, we decomposed single-cell RNA-seq data from HSCs that were followed by video microscopy to evaluate symmetric versus asymmetric distribution of lysosomes after divsion^50^. We found that in cells that had divided asymmetrically, the stem cell daughter more highly activated GRP18, compared to cells that had divided symmetrically (Figure 3E); all other GRPs did not differ significantly between asymmetric and symmetric divisions. In sum, these analyses suggest that GRP18 is associated with the propensity of HSCs to generate progenitors and divide asymmetrically.

### Co-regulation signatures improve transcriptome-based prediction of HSC functions

We then quantified GRP activity in a second clonal tracking experiment (“DARLIN”) that investigated clonal function in unperturbed hematopoiesis^51^. Here, clonal labels were induced by Cas9-mediating scarring in embryonic development, and both HSC transcriptomes and HSC output were determined in adulthood. We found strong associations between clonal activity/output and the GRPs related to cell cycle (GRP21 and GRP6), suggesting differences in the regulation of post-transplant and homeostatic stem cell output (Figure 3F).

To quantitatively determine how informative the single GRPs are of higher-order clone function, we applied machine learning (Figure 3G). We first determined how well HSC clonal function (activity and output, in the LARRY and DARLIN settings) can be predicted from HSC transcriptomes using black-box machine learning strategies. This resulted in an upper bound for how “explainable” clone function is from the transcriptome. We then compared the predictive performance of this approach to prediction based on a) the GRPs, b) gene lists commonly used in the field to characterize HSC function^4^, and c) co-expression programs identified by NMF. In all cases, we found that GRPs performed as well or better than gene lists or co-expression programs, suggesting a specific value of co-regulation signatures for predicting HSC functional variation (Figure 3G).

### GRPs are dysregulated in physiological aging

Together, decomposition analyses of clonal tracing experiments helped refine GRP annotation and demonstrated that co-regulation signatures can serve as an interpretable link between cell biological processes and HSC function. These results motivated a further exploration of GRPs in the context of human ageing and disease. We first used a large data set of blood stem- and progenitor cells in human aging^52^. Here, single-cell transcriptomes of CD34+ enriched blood cells had been collected from 148 donors aged 23-91. We applied decomposition to the “pseudobulk” gene expression of all cells annotated as HSC/MPP per donor, and found that one of the GRPs most strongly associated with aging was *inflammation response* (GRP3, Figure S8A), in line with literature^53^. Of note, inflammation response was low in young donors but highly variable in elderly donors, and on its own explained only 21% of age variation (Figure S8A,C). We additionally found significant associations between other programs and age, such as *lymphoid priming* (GRP14: lower in age, as expected^54^), and the *putative fetal HSC identity* signature (GRP13: decreased in age) (Figure S8B). A simple linear model of all GRPs explained 34% of variation in age on held-out test data (Figure S8C), modestly less than using full transcriptome information (39%). Collectively, these analyses illustrate how GRPs defined from Perturb-seq can be used to interpret gene expression data from human cohorts.

### A gene regulatory program predicts long-term survival in AML

Beyond applications in the context of healthy hematopoiesis, we attempted to use Perturb-seq GRPs to interpret and predict outcomes in AML. AML frequently has its origin in HSCs, and LSC properties are believed to impact long-term outcomes^55,56^. We made use of a single-cell RNA-seq atlas of diagnostic bone marrow aspirates from 184 AML patients, with annotation of response to first line therapy (for 133 patients treated with a similar regimen of intense chemotherapy), overall survival (**OS**, for all 184 patients), and relapse-free survival post allogeneic transplant (**RFS**, for 66 patients who received an allogeneic stem cell transplant)^57^. Importantly, the single-cell nature of the dataset allowed us to identify LSCs by transcriptomic similarity to healthy HSCs^58^. By applying decomposition to patient-level LSC gene expression profiles, we were able to associate GRPs with OS, RFS and first line response. Notably, higher cell cycle gene expression (*mitotic machinery*, GRP21) was associated with better outcomes in all categories, and GRP1 was strongly associated with better OS (Figure 4A).

In parallel work, we have found that first-line therapy response can be well predicted using various transcriptome-derived measures, including cell cycle, whereas relapse after allogeneic stem cell transplant is dominantly driven by microenvironmental, HSC-extrinsic features. By contrast, OS was difficult to predict from transcriptomic features^57^. Here, we surprisingly observed that in the context of OS, one particular GRP (GRP1) outperformed other transcriptomic measures, and served as a powerful predictor of survival in younger patients (Figure 4B,C). High levels of GRP1 activity were shared by long-term survivors with different mutational drivers (Figure 4D). Whilst GRP1 cannot trivially be mapped to a simple cell biological process, we observed that it was enriched for surface markers (*Sell*, *Selp*, *Met*, *Cd93*, *Itgb3*) with high expression in endosteal niches (Figure S9A-C), and we found a trend towards lower GRP1 activity in bone marrow derived LSCs from patients displaying extramedullary disease^59^ (myeloid sarcoma, Figure S9D). Together, these analyses suggest that GRP1 is associated both with long-term survival in AML, and with the ability of HSCs to interact with non-hematopoietic tissue.

### The “asymmetric division” GRP18 predicts response to Venetoclax

Unlike intensive chemotherapy, the BH3-mimetic Venetoclax is thought to specifically target LSCs by disrupting their energy metabolism^60,61^. We therefore asked if GRP activity could be used to predict Venetoclax response in a cohort of patients whose LSC transcriptomes had been profiled from flow-sorted GPR56+CD11b-CD64-cells^62^ (n=91 patients with data available for Ven/Aza response), where GPR56 is an LSC marker^63^ and double positive CD11b/CD64 mature myeloid cells were excluded. In line with the effect of Venetoclax on mitochondria, we found a significant association between higher activity of *mitochondrial activity* (GRP8) and Venetoclax response (Figure 4E). An even stronger association was identified with the *asymmetric division* GRP18. GRP18 predicted Venetoclax response better than a clinically used genetic signature^64^ in 10-fold cross validation, and performed slightly inferior to a recently developed flow cytometric score^65^ (Figure 4F,G). It substantially outperformed NMF-derived co-expression signatures. This result suggests a potential value of Perturb-seq GRPs for interpreting therapy response data, and a link between asymmetric divisions and vulnerability to Venetoclax. By design, our approach identified candidate genetic regulators of all AML-related programs (Figure S9E-G).

## Discussion

The inference of gene functions and regulatory interactions by measuring the phenotypes of perturbations is a core tenet of genetics. Perturb-seq – the simultaneous profiling of hundreds to thousands of gene perturbations with a single-cell transcriptomic read-out – extends this paradigm to a systems-level scale, but data in disease-relevant primary cell models remains scarce.

In this study, we performed Perturb-seq in stem cell expansion cultures established from primary mouse bone marrow, profiling transcriptome-wide changes induced by over 500 perturbations in the stem and progenitor cell population. This focus distinguishes our work from previous studies applying Perturb-seq to investigate cell state changes during hematopoietic differentiation^5,11^. Perturb-seq in this model poses analytical and technical hurdles related to effects of perturbations on differentiation state, as well as the potential for false positive perturbation effects due to clone-associated heterogeneity. Experimental design choices such as the inclusion of 3 guide RNAs per target gene to evaluate consistency between guide effects, or the use of >100 non-targeting control guides, as well as bespoke analytical tools, were both important for mitigating these problems. Future screens could directly resolve the contribution of clonal heterogeneity through incorporating unique clonal barcodes in the experimental design^32^.

By using neighborhood correction to mask effects of knockdown on cellular differentiation state, we emphasized local perturbation effects on gene expression within cell states. These perturbation signatures facilitated the grouping of perturbations by interacting genes within protein complexes, and provided the basis for the derivation of 19 genetic programs of co-regulation. Most GRPs demonstrated strong correspondence to cell biological processes and were relatively consistent across HSC and early progenitor cell states. Together, these observations suggest broad genetic co-regulation of cellular pathways, e.g. components of the Nfkb signaling pathway, or translation initiation factors.

We demonstrated the decomposition of external HSPC transcriptome data into these programs, an approach that we propose to be broadly useful in many contexts. GRPs were associated with functional variation of HSCs and performed as well or better than co-expression programs and expert-curated signatures for machine learning-based prediction. Furthermore, when decomposition was applied to human data sets, GRPs were predictors of stem-cell-related clinical outcomes such as long-term survival or Venetoclax response in retrospective AML cohorts. These results suggest that co-regulation programs carry important and previously undescribed relevance for higher-order biological function.

We propose that GRPs, inferred from genetic perturbation data (Perturb-seq), readily associate with cell biological processes and stem cell functions, and can serve as a reusable vocabulary for interpreting HSC/MPP transcriptomes. This comprehensive approach allowed us, for example, to map asymmetric cell division, stem cell output post-transplant, and venetoclax sensitivity to a single program, and to implicate a Myc-Myb axis in its regulation. This inherent feature of the GRPs – that their genetic regulators are known – opens the possibility of identifying and prioritizing regulators that modulate HSPC states.

In sum, our work introduces programs of co-regulated genes, their regulation, their relationship to cell biological processes, and their association with stem-cell-associated functional and disease phenotypes (Table 1).

### Limitations

Our study reports data for the knockdown of 520 genes. Whilst these genes were prioritized to include many important regulators of hematopoiesis, we inevitably missed some relevant regulators. Moreover, even for included genes, some perturbation effects may be obscured by limited cellular coverage, inefficient repression by CRISPRi, or clonal heterogeneity dominating over weak perturbations. Notably, our analytical approach to derive GRPs involved subsetting the data to perturbations with relatively stronger effects, which in practice likely improved GRP inference by limiting variability introduced by cells with weaker perturbations. While we showed that GRPs are relatively consistent across HSC and MPP states, a larger screen of pure HSCs (e.g. LSK EPCR+ cells) could help to further refine HSC-relevant programs, but poses significant hurdles regarding availability of material and screen size.

## Data availability

Raw files of Perturb-seq data generated in this study are available in the Gene Expression Omnibus (GEO) under accession number GSE320250. Seurat objects containing the raw, normalized, and neighborhood-transformed single-cell expression matrices are available at https://doi.org/10.6084/m9.figshare.31276981. Tables of differential expression results for all perturbations are available at https://doi.org/10.6084/m9.figshare.31277494. Tables of differential abundance results are available at https://doi.org/10.6084/m9.figshare.32531301.

## Code availability

An R package for HSPC transcriptome decomposition is available at https://github.com/veltenlab/hsc19. Package vignettes contain code to reproduce most figures of this manuscript.

## Acknowledgements

We thank Adam Wilkinson for sharing protocols and David Lara-Astiaso, as well as all members of the Velten lab, for feedback on the manuscript. We thank the CRG facilities for Genomics and Flow Cytometry, as well as the PRBB animal house, for experimental support. This project has received funding from the European Union’s Horizon Europe under the grant agreement no. 101041399 (ERC-StG AI4SYN to L.V.). The authors acknowledge support of the Spanish Ministry of Science and Innovation to the EMBL partnership, the Centro de Excelencia Severo Ochoa and the CERCA Programme / Generalitat de Catalunya.

## Author contributions

JB and LV conceived of the study, analyzed the data and wrote the manuscript. JB performed experiments, with support by ABM. JSS contributed to experimental design. All other authors contributed to the analysis of AML data. All authors commented on the manuscript.

## Overview of materials appended below

- Table 1
- Figures 1-4 and legends
- References
- Methods
- Supplementary Figures and legends, Supplementary Table legends
- Supplementary Note

## Methods

### Experimental Methods

#### Selection of genes in CRISPRi Perturb-seq

From literature^28^ and power calculations on pilot experiments, we aimed for an average of 200 cells per gene targeted by CRISPRi, equating to ∼500 genes target genes in an experiment of approximately 100,000 cells. We selected 520 genes by a ranking system which assigned each gene in the mouse genome a score out of 100 based on relative weighting of the following criteria:

1. Expression parameters evaluated from existing ex vivo mouse hematopoietic stem-and progenitor culture scRNA-seq data (40/100)
2. Gene Ontology (GO) annotation with terms related to transcription factors or chromatin-associated cofactors (20/100)
3. Count of entries in PubMed containing the gene name in the context of hematopoiesis or similar keywords (20/100)
4. Overlap with transcription factors included in ^67,68^ (20/100)

This ranking table is provided in full as Table S1. Of the top-ranked genes, 20 were prioritized for inclusion at 5-fold in the final guide RNA library.

#### Guide RNA library design and cloning into CROP-seq-Puro-eGFP

Guide sequences for 3 CRISPRi guide RNAs per target gene were designed with the CRISPick tool (Broad Institute) using ‘Library Mode’ to additionally select 120 control guides – 60 with no sequence homology in the mouse genome (‘NO-SITE’) and 60 targeting only one intergenic region of the genome (‘ONE-INTERGENIC’). The library was ordered as an oligo pool from Twist Biosciences with the target-specific spacer sequences flanked by universal amplification handles and Esp3I motifs as follows:

5’-*GAAGTGCCATTCCGCCTGACCTCGTCTCACACCG*/spacer/*GTTTCGAGACGAGGCTAGGTGGAGGCTCAGTG*-3’ 5-fold coverage of prioritized guides (60 targeting + 20 control) was achieved by repeating lines in the spreadsheet for ordering.

The guide RNA library was cloned as a pool into a modified^28^ version of the CROP-seq vector^10^ that included the Chen F+E tracr variant^69^ and a Puro-eGFP marker for selection. Briefly, the lyophilized Twist library was first resuspended to 50 fmol/µl with 10 mM Tris pH8.0. Then, PCR amplification for 8 cycles was performed using biotinylated primers/5Biosg/GAAGTGCCATTCCGCCTGACCT (forward) and /5Biosg/CACTGAGCCTCCACCTAGCCT (reverse) and NEBNext High-Fidelity 2x PCR Master Mix: 98°C for 30 s, 8 cycles of [98°C for 10 s, 61°C for 15 s, 72 °C for 10 s] and a final extension of 72°C for 60 s. The Twist amplicon was digested with Esp3I-HF at 37°C for 2 hours in T4 Ligation buffer (with ATP) without heat inactivation. To remove biotinylated primers, the digestion mix was incubated for 10 minutes at room temperature with Streptavidin T1 Dynabeads and the supernatant containing the insert of the guide RNA library was kept. Separately, the CROP-seq-Puro-eGFP backbone vector was digested with Esp3I-HF and purified by gel electrophoresis. The guide library insert was ligated into the digested backbone at a molar ratio of 20:1 insert:vector (∼5:200 ng insert:backbone) with T4 DNA Ligase. Ligation was performed at 21°C for 80 minutes then 65°C for 10 minutes and the final ligation product containing assembled libraries was purified with DNA Zymo Clean and Concentrator -5 kit. This was transformed into Lucigen Endura electrocompetant bacteria and plated to multiple LB+Ampicillin agar plates. An estimated 1.8x10^5^ bacterial colonies were counted (coverage of ∼100 per guide) and scraped from the plates for DNA extraction using PureLink Plasmid Maxiprep kit (Invitrogen).

Successful cloning was confirmed and the distribution the guide RNAs within final Perturb-seq library (Figure S1A) quantified by PCR amplification over the guide RNA sequence using primers with Illumina sequencing handles following by next-generation sequencing on an Illumina iSeq machine.

#### Lentivirus production

HEK293FT cells used for lentivirus production were cultured with DMEM high glucose (Thermo Fisher 11965092) supplemented with 10% FBS (A5256701), 1% Glutamax (35050061), 1 mM sodium pyruvate (11360070) and MEM non-essential amino acids (11140035). Cells at 75% confluency were transfected with plasmids psPAX2 (1.3 pM, Addgene #12260), pMD2.G (0.7 pM, Addgene #12259) and CROP-seq-Puro-eGFP with the CRISPRi guide RNA library (1.64 pM). Transfection was performed with Lipofectamine3000 for 6 hours, then the transfection mix was replaced with fresh media. Cell culture supernatant containing produced virus was collected on day 3 post-transfection. Media was cleared through a 0.45-µm filter, treated for 20 minutes (37°C) with 1 mg/ml DNAse and 1mM MgCl_2_, and precipitated with sucrose cushion ultracentrifugation (6000g for 90 min at 4°C)^70^. Concentrated lentivirus pellets were washed once with cold PBS and stored after snap-freezing as aliquots in PBS at -70 °C. A separate round of lentivirus production was performed ahead of the second replicate experiment.

#### CRISPRi mouse lines

We generated experimental animals with constitutive expression of the CRISPRi repressor in hematopoietic lineages by a cross between Rosa26-LSL-dCas9-KRAB^29^ (obtained from the Jackson laboratories, strain ID #033066) and Vav1-Cre^30^ (obtained from the lab of Thomas Graf, Jackson strain ID #035670) strains. F1 offspring were genotyped by PCR as recommended in Jax #033066 and #035670 to confirm animals heterozygous positive for both genotypes. All procedures involving animals adhered to the pertinent regulations and guidelines of the European Union, Spain, and Catalonia. Approval and oversight for all protocols and strains of mice were granted by the Institutional Review Board and the Institutional Animal Care and Use Committee at PRBB, under protocol 11733.

#### HSC expansion cultures

For the main Perturb-seq screen (Replicate 1), ex vivo HSC expansion cultures were initiated from 5 adult male mice at 6 months old (Replicate 2 used cells collected from different animals: 3 females, 2 males). Mice were euthanized by cervical dislocation and the bones from limbs were extracted and cleaned from connective and muscle tissue. Bones were crushed in PBS to release marrow cells, filtered through a 40 µm strainer, and pelleted by centrifugation at 400 g for 5 min at 4°C. Red blood cells were lysed with 1X ACK Lysis buffer and remaining cells were enriched for HSCs by two rounds of selection using MACS columns: first lineage depletion using a mouse Lineage Cell Depletion Kit (Miltenyi Biotec 130-110-470) (keep the flowthrough), then c-Kit enrichment with CD117 MicroBeads (Miltenyi Biotec 130-091-224). Of note, this procedure enriches but is not completely selective for Lin-cKit+ cells, as measured by flow cytometry.

Further enrichment and expansion of HSCs was achieved by the culture conditions and media composition reported in detail in ref. ^71^. 0.75x10^6^ cells were initially plated to each of 24-well CellBIND plates (Corning 3337) and cells were incubated in humid hypoxic conditions (5% O_2_, 5% CO_2_, 37 °C). Media changes, each time with cytokines freshly added to media from frozen aliquots, were performed carefully every 2 days (see Video 1 from ref. ^71^). From the stage where cell growth inwards from the edges of the plates begins to cover (semi-adherently) the surfaces of the wells (day ∼10) to day ∼50, cultures remained in state of stable proliferation and were passaged to new 24-well plates with a 1:2 dilution every 4 days. Cultures were monitored by flow cytometry and contained approximately 60% Lineage-cKit+ Sca-1+ cells. For Replicate 2, cultures were restored from vials frozen in CellBANKER 2 media.

#### Flow cytometry

The health and status of the cultures was monitored by staining a fraction of the cultures (∼1-5x10^5^ cells) with fluorescently-labelled antibodies: FITC anti-mouse Lineage Cocktail (Biolegend 78022), BV605 anti-mouse cKit/CD117 (Biolegend 105847, clone 2B8), BV785 anti-mouse Ly6A/E(Sca-1) (Bio Legend 108139, clone D7), APC anti-CD201 (EPCR) (Biolegend 141505, clone RCR-16). Cell sorting was performed using BD FACSAria II or BD Influx instruments.

#### Lentiviral infection

12 tubes of 1ml were prepared for infection by mixing 1.8x10^6^ cells with lentivirus in HSC media supplemented with Protamine Sulphate (20ug/ml). Each infection mix was plated to 10 wells of 96-well Fibronectin plates. After 24 hours, infected cells were washed with PBS and re-plated to fresh CellBIND plates. 7 days following initial infection, live GFP+ cells (2.5x10^5^ cells, 1.15% of live) harbouring lentiviral integrations were collected by FACS and fixed immediately using the Evercode Fixation kit from Parse Biosciences (ECFC3300) and stored at -70°C.

#### Single-cell RNA-seq by split-pool combinatorial barcoding

All fixed GFP+ cells were processed for single-cell RNA-seq using an Evercode Whole Transcriptome v3 kit (ECWT3300, Parse Biosciences). An Evercode Mini kit (ECWT3100) was used for replicate 2. The CRISPR Detect accessory kit (CRS1010) was used to specifically amplify next-generation sequencing sublibraries of guide RNAs expressed via the CROP-seq transcript. All whole-transcriptome and CRISPR guide sublibraries were sequenced on an Illumina NovaSeq S2 machine according to the recommendations from Parse Biosciences: read cycles of 64-8-8-58, ∼30,000 reads per cell, and 5-10% PhiX spike-in.

### Computational Methods

#### Quantification of guide RNA composition in the CROP-seq vector

The ‘guide-counter’ tool from Fulcrum genomics (https://github.com/fulcrumgenomics/guide-counter) was used to quantify the composition of guide RNAs within the library cloned into the CROP-seq-Puro-eGFP vector. Paired fastq reads were mapped to a dictionary of spacer sequences provided for each guide RNA, allowing by default settings for 1 mismatch. The resulting table of counts of each guide in the vector was used for Figure S1A. For the essentiality analysis (Figure S1G), the proportions of each guide in the CROP-seq vector (reads) were compared to the proportions of cells containing each guide in the final Perturb-seq data. We used edgeR’s negative binomial GLM framework^72^ to test for differences in guide proportions, with the 3 guides targeting each gene considered as replicates in a paired design. Significance was calculated using likelihood ratio tests and adjusted for multiple testing (Benjamini-Hochberg correction).

#### Mapping of NGS libraries in CRISPRi Perturb-seq data

Fastq files of whole-transcriptome sublibraries were processed using the *split-pipe* pipeline provided by Parse Biosciences. In this pipeline, reads are first assigned to single cells according to the combinatorial index recorded in Read1 then mapped to a reference genome and assigned to genes. For this data set, the mouse reference genome was built with ‘*split-pipe --mode mkref*’ from Ensembl (release 112) GRCm39 primary assembly and genome annotation. Data from 8 sublibraries containing single-cell transcriptomes were combined with ‘*split-pipe --mode comb’* and the gene expression counts matrix (cells x genes) was made into a Seurat object in R for further processing.

Libraries of CRISPR guide RNAs were initially mapped with minimal read and transcript thresholds (parameters ‘*split-pipe --crispr --crsp_read_thresh 3 --crsp_tscp_thresh 1’*). A transcript threshold of 2 was applied downstream in R for guide assignment to single cells within the Seurat data object. We noticed that more cells than expected contained multiple guide RNAs targeting the same gene. We identified this as caused by PCR/mapping artifacts from guide RNAs in our library which differed from each other by a shift of only 1 or 2 bp. These cases were collapsed to the guide with more reads by a custom R function. We additionally used the crispat package^73^ to evaluate other methods for guide assignment strategies. No other strategies outperformed the simple two-transcript threshold, likely because guide RNA detection from CROP-seq yields fewer transcripts per cell in primary cell models compared to cell lines.

#### Perturb-seq analysis

The combined data from the CRISPRi Perturb-seq experiment (Replicate 1) was filtered with Seurat^74^ to remove outlier cells on the basis of transcript count, number of detected genes, or percentage of mitochondrial reads. This resulted in a single-cell Seurat object of 111,003 single cells. Transcriptomes were normalized by SCTransform^75^ and processed to generate the UMAP^76^ embedding shown in Figure 1 using ‘*RunPCA*’ and ‘*RunUMAP(dims=1:30)*’. To annotate cell states, we first performed Leiden clustering^77^ on the shared nearest-neighbor graph computed from the first 30 principal components. Clusters were annotated based on marker gene expression. Cell cycling scoring was performed using mouse orthologs of the curated S-phase and G2/M-phase gene sets^78^ provided in Seurat. The annotated Seurat data object is available on Figshare.

On-target expression changes (Figure S1F) were calculated using ‘*FindMarkers’*, comparing cells with the tested guide or gene versus cells containing non-targeting guides. Only cells assigned a single guide RNA identity were used. Differential abundance analysis across different cell states (Figure 1E, Figure S3A,B) was performed on contingency tables constructed from cells containing single control guides. Fisher’s exact tests were used to test for enrichment of counts of cells with a particular perturbation compared to cells containing non-targeting control guides.

#### Differential expression analysis using SCEPTRE

We used the R package SCEPTRE^33^ for Perturb-seq differential expression analysis. A more detailed explanation of the methodological considerations of this analysis is described in Supplementary Note.

#### Neighborhood transformation of perturbed transcriptomes

First, we compiled a set of response genes (n = 2875) by combining the 1000 most variable genes in the data with the 10 most significantly differentially expressed genes from each perturbation. Using these response genes, neighborhood transformation was performed on the SCT assay using the ‘*CalcPerturbSig*’ function from the Mixscale package^34^ within Seurat. This transformed the transcriptome (log1p) of each perturbed cell by subtracting its expression from the average expression of its 20 nearest neighbours control cells in PCA space (40 dimensions): Expression(control neighbours) – Expression(perturbed cell). Notably, this flips the signs, so that a repressed gene (e.g. the target gene of a guide), which has lower expression in the perturbed cell than its neighbours, becomes transformed into a positive value. Gene expression vectors after this transformation (aka the ‘PRTB matrix’) were scaled and centred, with technical covariates related to library size (number of transcripts and genes per cell) regressed out.

We used the first 15 principal components from the PRTB matrix to generate UMAPs following neighborhood transformation (Figure S4A,B) after elbow plots and JackStraw analysis demonstrated that additional PCs contributed little biological variance. This reflects that neighborhood correction removes much of the variation in the data unrelated to perturbations.

#### Correlation of “pseudobulk” perturbation signatures

We subsetted to only cells containing one guide RNA and calculated the average expression per perturbation (guide) of the n=2875 response genes defined above, from both untransformed (SCT) and neighborhood-transformed matrixes (PRTB). Pairwise correlations (Pearson R values) were calculated from these “pseudobulks” (Figure S4C).

#### Filtering for consistent perturbations

We derived the ‘Same-Gene-Similarity’ (SGS) score as a measure of consistency in transcriptome signatures of perturbations between guides targeting the same gene. This was defined for each perturbation from the pairwise distance matrix (1 – Pearson R) as:

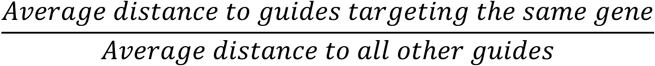

An SGS score of 0 represents perfect similarity between perturbation signatures while a score of ∼1 represents no similarity. Simulations were performed to define baseline distributions of SGS scores using randomly-grouped triplets of control guides, and perturbations with SGS scores lower than the 5^th^ percentile were classified as “consistent”. This process was performed at both the level of guide perturbations and perturbed genes (average of SGS scores) (Figure S4E). The pairwise distance matrix was then filtered by these criteria to only include consistent perturbations for generating the embedding in Figure 1G.

#### Cell state-specific perturbation signatures

To compare perturbation signatures separately across cell states, we made five subsets of the Perturb-seq Seurat object, which had already subsetted for consistent perturbations. These subsets were: HSC/MPP, MyP and MegEry, based on the cell state annotations described above (Figure 1C); Hlf+, defined as any cells with counts for *Hlf*; and Hlf+ Mecom+, a smaller “transcriptome-defined HSC” subset defined as cells with >2 counts for both *Hlf* and *Mecom*. We then made perturbation pseudobulks from the PRTB assay for each subset and calculated pairwise Pearson correlations between the same perturbations across different cell states.

#### Embedding and clustering of co-regulating perturbations

We employed a two-step dimensionality reduction workflow on the distance matrix of consistent perturbations. We first used spectral embedding^79^ to capture the manifold structure, then t-SNE for two-dimensional visualization of local neighborhoods. We used parameters of “*n_components = 40, n_neighbors = 3, affinity = ‘precomputed_nearest_neighbors’, eigen_solver = ‘arpack’*” for ‘SpectralEmbedding’ and “*perplexity = 5*” for ‘TSNE’, both functions within *scikit-learn*. These parameters were determined by a sweep aimed to minimize Euclidean tSNE distance between guides targeting the same gene.

Clustering of co-regulating perturbations was performed using HDBSCAN^80^ on the spectral embedding with the following parameters: “*metric = ‘euclidean’, min_cluster_size = 2, min_samples = 1, cluster_selection_method = ‘eom’, cluster_selection_epsilon = 0.13, prediction_data = True*”. A parameter sweep was performed to determine HDBSCAN parameters to maximize grouping of guide perturbations targeting the same gene. After the primary clustering step, perturbation points initially classified as ‘noise’ by HDSCAN (ie assigned to cluster ‘-1’) were added to their nearest clusters, and some closely-related clusters were merged. A table of the perturbations (guide RNAs) included in each of the final clusters is provided as Table S2. We also provide a vignette to reproduce the embedding (see code availability statement).

#### Guided Sparse Factor Analysis (GSFA)

GSFA is a Bayesian statistical model that combines factor analysis with joint estimation of effects of perturbations on factors. The mathematical basis of GSFA is detailed in ref. ^41^. Practically, the package requires as input two matrices; a scaled gene expression matrix and a binary matrix of perturbation identity from same cells. Two key outputs are the W matrix, which defines latent factors in terms of weights of genes loaded to each factor, and the β matrix, which quantitatively associates each latent factor with perturbations. The W and β matrix are also paired with matrices which quantify the Bayesian PIPs (Posterior Inclusion Probabilities) for each entry in these matrices; loadings or perturbation associations with a low PIP are effectively negligible.

Our application of GSFA differs from Zhou et al in the use of neighborhood-transformed transcriptomes (i.e. PRTB matrix) as the scaled gene expression input. We extracted input matrices for GSFA from cells in the CRISPRi Perturb-seq data object carrying single guides targeting genes defined as ‘consistent’ by the SGS score criterium (117 of 520 genes, 23,515 perturbed cells). We then performed GSFA for 3000 iterations of Gibbs Sampling and used the final 2000 iterations to obtain the posterior sample estimates. Other parameters of our implementation were “*K = 21, prior_type = ‘mixture_normal’, init.method = ‘random’*”.

The choice of K = 21 latent factors was informed by the approach recommended in Supplementary Note 4 of Zhou et al. We performed GSFA at variable Ks, initially in steps of 5 (from 5 to 35), and then across all values of K in a critical range between 17 and 23. Calculations of the proportion of variance explained (PVE) of the starting PRTB matrix demonstrated saturation behaviour, but with relatively gradual saturation (Figure S6A). We eventually chose K=21 as a trade-off between increasing PVE and drawbacks of long compute times with greater K (non-linear) and reduced factor interpretability. Factors (defined by their W matrices) are generally robust to small changes in K parameter, with Pearson correlations between W matrices showing correspondence between factors (Figure S6B). The same parameters were also used to run GSFA on the cell state-specific subsets of the data described above.

The output matrices from the GSFA model fit used for downstream analyses are provided within our R package.

#### Non-negative Matrix Factorization (NMF)

We used the GeneNMF package^81^ to infer co-expression programs from the untransformed gene expression profiles in our Perturb-seq data. Specifically, we used the SCT matrix subsetted to only control cells and the same genes as the PRTB matrix. We ran *multiNMF* across subsets of the data, corresponding in practice to sublibraries in the Parse Evercode kit, and across k values from 5 to 16. We then used *getMetaPrograms* to summarize the results as k=15 metaprograms. These parameter choices were informed by tutorials on the GeneMNF GitHub: https://github.com/carmonalab/GeneNMF. Metaprograms were converted to the same data format as the W matrix from GSFA, namely sparse lists of genes with loading weights (median 56 genes per metaprogram) for comparison to GSFA factors by Pearson correlation.

#### Annotation of latent factors as gene regulatory programs (GRPs)

For downstream annotation and analysis of factors, we enforced an extra layer of sparsity on the GSFA gene loading matrix, *W*, by applying a posterior inclusion probability (PIP) filter of > 0.8 (greater than 0.8 likelihood of a gene loading to a factor). We also transformed *W*′matrices so that the majority of gene loadings for each factors carried positive weights. These sparse and sign-aligned factors aided factor annotation by allowing programs to be more consistently interpretated as positively associated with any associated biological process. Factor annotation was performed with the lists of genes both positively and negatively loaded to each factor. Our approach is summarized in Table 1 and combined 1) manual inspection and literature research of the lists of genes loaded to each factor (Table S3), 2) enrichment of gene lists in functional databases such as GO, KEGG or TF-targets, and 3) large-language model (GPT5.1) annotation. This approach was pioneered and benchmarked in ref. ^42^; we here used a more recent LLM (GPT5.1), and specifically the prompts:

> *I have provided you with files, named according to Factor_X, where X is a number between 1 and 21. Each file contains a list of mouse genes. For each factor, there is a list of top genes which are positively associated with the factor. And a list of bottom genes which are negatively associated with the factor. For each factor (pair of top and bottom files), ask: What do these genes have in common? And: what might their common role in hematopoietic stem cells be? Give a biological name for each factor (eg. translation initiation), and also give a value on how confident you are in this annotation.*
>
> *Can you repeat the same analysis, but keeping in mind that genes are ranked in the lists from most important to least important for the factors*.

GRP13 and GRP16 were labelled as ‘putative’ due to low confidence scores from LLM annotation and limited other associations.

#### Effects on perturbations on GRPs

A filter of PIP > 0.8 (from the ‘Gamma’ PIP matrix) was also applied to perturbation-factor effects (β matrix). Functions for plotting perturbation effects on factors were written in R. These often took inspiration from code shared with the original GSFA (xinhe-lab.github.io/GSFA_paper/), and are available on our GitHub page (see code availability statement).

#### Preprocessing of external transcriptome data sets

To link GRP activity to HSC function, we first collected published data sets that linked HSC gene expression to different functional outcomes.

1. *Human Ageing.* We downloaded single-cell gene expression data^52^ from https://cellxgene.cziscience.com/collections/5542eeb0-96ef-4ab9-95ea-eb6abc178461 and subsetted to all cells annotated as “HSC/MPP”. and subsetted to all cells annotated as “HSC/MPP”. Subsequently, we summed raw read counts per individual.
2. *Clones post-transplant (LARRY).* We downloaded clonally barcoded single-cell gene expression data of Lineage-negative, c-Kit-positive cells from five different experimental animals^48^ from https://doi.org/10.6084/m9.figshare.24260743 and subsetted to all cells annotated as “HSC/MPP1”. Subsequently, we summed raw read counts per clone.
3. *Clones in unperturbed hematopoiesis (DARLIN).* We downloaded clonally barcoded single-cell gene expression data^51^ and associated scripts from https://doi.org/10.5281/zenodo.11929508. We then used the script DARLIN_tutorial/Single-cell-tutorial-Part_2_downstream_analysis.ipynb to prepare a data object containing both clone and gene expression information. We subsetted to all cells annotated as “HSCs”, and summed raw read counts per clone.
4. *Symmetric vs. asymmetric divsions.* We used a data set of HSCs that had been followed by live cell imaging to determine post-mitotic distribution of lysosomes^50^ prior to single cell RNA-seq. Single-cell RNA-seq count matrices were obtained from GEO (GSE167317). Divisions were classified as symmetric or asymmetric based on the ratios of the “LysoBriteOMA_norm” metadata column (see figure 3E).
5. *Human AML – survival and chemotherapy response.* We used a single-cell RNA-seq dataset of 184 AML patients, selected to represent most genetically defined (WHO) groups^57^. This dataset had been annotated by projection on a healthy reference^58^, to identify the most similar healthy cell type / differentiation state for each cell. We subsetted to cells corresponding to stem and progenitor cells (i.e. up to the promyelocyte stage, and excluding cells resembling mature monocytes, and other mature myeloid, erythroid and lymphoid cells), and summed raw read counts per patient. Detailed outcome information was available, including overall survival, response to first-line chemotherapy (for n=133 patients treated with intense 7+3 chemotherapy), if allogeneic treatment was used before or after relapse, and time until relapse. Importantly, the dataset also provided relevant potential predictors of outcomes that we compared to the GRPs: Genetic classification and clinical metadata (WHO classification, ELN-22 risk score, patient age), expert-curated transcriptome derived scores (LSC-17 score^21^, “pseudotime” of differentiation arrest^58^, cell-cycle-related scores), and information on non-leukemic cells (e.g. immune cell composition).
6. *Human AML – Venetoclax response.* We used bulk RNA-seq dataset of sorted LSCs (GPR56^+^CD11b^-^CD64^-^) from 91 AML patients treated with Venetoclax+Azacitdine, with response information available. Data were downloaded from https://www.ebi.ac.uk/biostudies/ArrayExpress/studies/E-MTAB-14142?key=c1957a18-a2d0-49cf-bd15-d66d2dcf98b0. This dataset also provided classification of patients by a 3-gene score currently used to predict Ven+Aza response^64^, as well as other clinical metadata.

For each of these datasets, we accordingly obtained raw gene expression count matrices of HSCs aggregated at the level of individuals or clones (or single cells, in the case of the symmetric/asymmetric cell division data). Subsequently, we applied a coherent pre-processing workflow using the Seurat package. Specifically, data were log-normalized, highly variable genes were identified and principal component analysis was performed for downstream machine learning tasks. For comparison to the GRPs, we also used the Seurat AddModuleScore function to calculate expression scores of gene lists commonly used in the field, as curated in Table S2 of ref. ^4^: Rodriguez-Fraticelli et al. Low output, High output, Mk bias, and serial engraftment HSC signatures^4^, Rodriguez-Fraticelli et al. HSC clusters 1-4, Giladi et al. Stem-score^82^, Lauridsen et al. Retinoic Acid Report-dim signature^83^, Wilson et al. signatures linking HSC phenotypes and function^84^, Cabezas-Wallscheid et al. dormant vs. active HSC signatures^49^ and Pietras et al. HSC signature^85^. Finally we computed cell cycle signatures^78^.

#### Decomposition of HSPC transcriptome data into GRPs

To decompose arbitrary HSPC transcriptome data into the GRPs, we treated log-normalized gene expression data as a linear combination of the sparse GRP loading matrix *W*′, defined above. After subset solved by ordinary least squares *x_i_* = *W*′*ζ_i_*

where *x_i_* is the normalized gene expression vector corresponding to sample *i*, and *ζ_i_* is the estimated activity of each GRP in the sample. Unlike in cell type decomposition (e.g. CIBERSORT^86^), where the “signature matrix” is computed from measured gene expression profiles, *W*′ stems from a Bayesian inference procedure and is less noisy; we therefore did not observe improved performance when using support vector regression (as in CIBERSORT) instead of ordinary least squares.

To evaluate statistical association between inferred GRP activities and outcomes of interest, we used a correlation test for continuous outcomes (e.g. age), a Wilcoxon test for binary outcomes (e.g. Venetoclax response) and a Cox regression for survival outcomes.

#### Evaluation of the predictive value of GRPs

We aimed to evaluate the predictive value of a GRP on an outcome (such as overall survival in AML), to compare the predictive value of GRPs to other transcriptome derived scores^4^, other predictors (such as genetic classification), and to estimate the amount of predictive information contained in the whole transcriptome. To that end, we set up a 10-fold cross validation scheme, i.e. we split the data into 10 equal-sized folds, nine of which were used for model training (train set) and one for model evaluation (test set). Within the train set, we performed the following steps:

1. We identified the single GRP with the highest correlation with the outcome, using linear regression, generalized linear regression, or cox regression, depending on the type of outcome. We then used that GRP for predicting the outcome on the test set. By selecting a single GRP at training, we thereby ensure that there is no benefit of testing multiple GRPs, e.g. when comparing to a single other predictor.
2. From a list of pre-defined transcriptomic gene sets commonly used in the field^4^ (see above, *Preprocessing of external transcriptome data sets*), we computed expression scores, equally picked the best using the train set, and used it for predicting the outcome on the test set.
3. In the context of AML, we additionally trained models to predict outcome from clinical metadata (such as patient age, ELN risk scores, WHO classification or the LSC17 score). Each of these models were trained separately and separately evaluated on the test data.
4. Finally, we trained several machine learning models using all transcriptome data and evaluated each model on the test data. Specifically, we used principal component regression, principal component LASSO, LASSO and regular regression of differentially expressed genes (e.g. genes DE between long-term survivors and short-term survivors in AML, with DE performed on the train set only), and LASSO regression on pre-defined gene sets^4^. Of these models, we include the model with the best performance as an upper bound (labeled “full transcriptome (ML)” in figures 3F and 4B,G).

We then repeated the same procedure with all 10 folds as test data and compute the estimate performance (for numerical outcomes: coefficient of determination / R^2^; for binary outcomes: area under the curve / AUC; for survival outcomes: C-index) on across all test sets. Finally, we repeated this procedure 10 times with different train/test splits, to obtain a confidence interval on the performance metric.

## Supplementary Figures and legends

**Figure S1.**
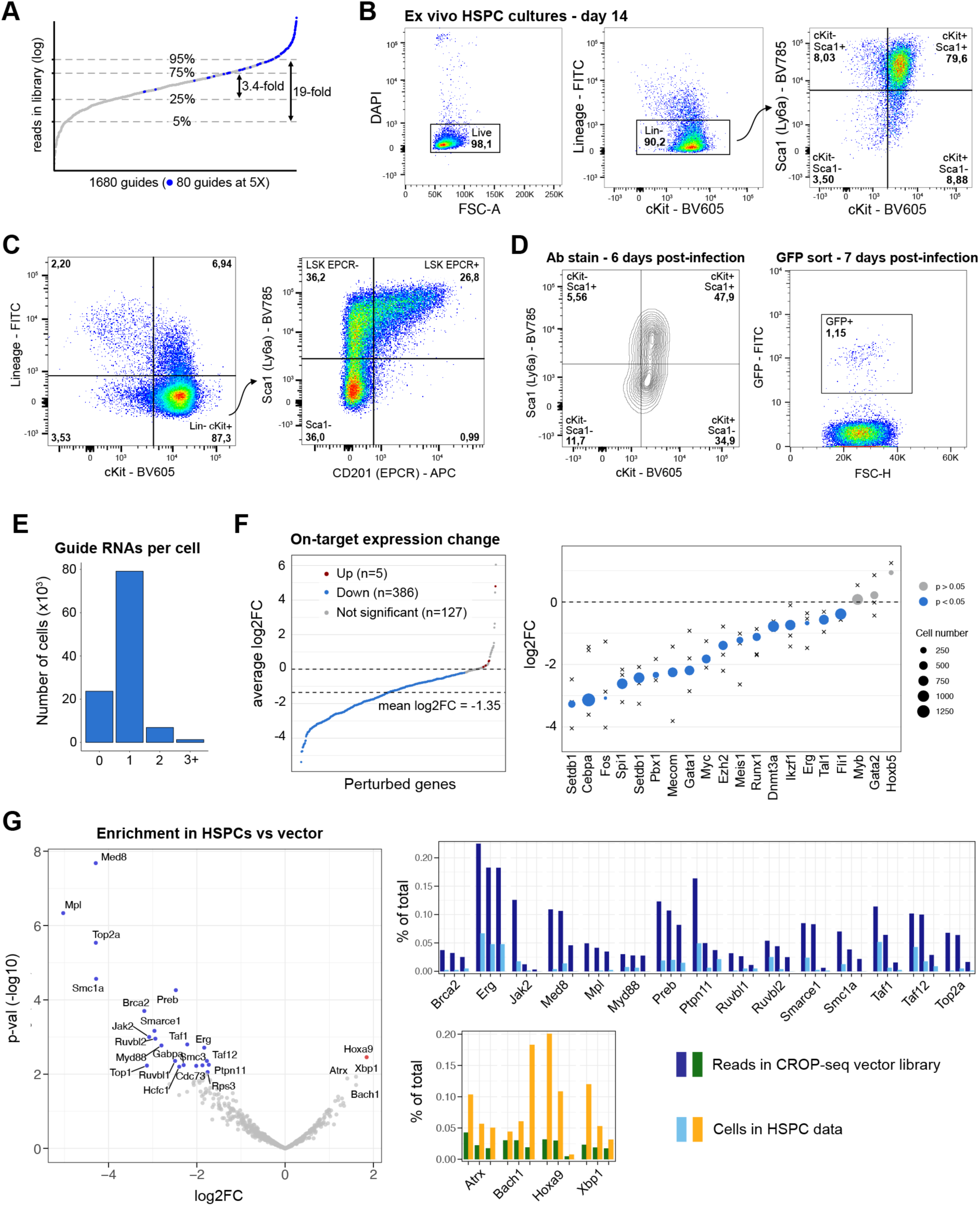
Successful implementation of Perturb-seq in HSC expansion cultures, related to Figure 1. **A.** Proportions of each guide RNAs within the library cloned into the CROP-seq-GFP vector, measured from next-generation sequencing reads. Blue points show guide RNAs that were intentionally included at 5-fold representation in the library (60 targeting, 20 control) and are excluded from fold change calculations of 25-75% and 5-95% ranges. **B.** Antibody staining flow cytometry characterization of the cultures used for the Perturb-seq experiment, a few days before infection. Fluorescent antibodies used are provided in Methods. **C.** Antibody stain demonstrating the typical fractions of immunophenotypically defined HSCs (Lin-cKit+Sca1+EPCR+) in ex vivo HSPC cultures in our hands. Note this panel shows data from a later experiment rather than the exact culture used for this Perturb-seq experiment. **D.** Left: Antibody staining flow cytometry characterization of infected cells a day before collection for scRNA-seq. Right: FACS gate used to sort GFP+ cells on day of collection. **E.** Bar chart quantifying how many unique guides are detected in each cell profiled in Perturb-seq. **F.** Quantification of on-target differential expression changes in Perturb-seq. Left: Plot of all targeted genes ordered by magnitude of fold change between perturbed cells and controls. Significance was calculated by the Wilcoxon test implemented by Seurat’s *FindMarkers* and corrected for multiple-testing (Benjamini-Hochberg). Note Cdx4 and Pou5f1 are missing due to insufficient counts. Right: Plot of genes included at 5-fold (x) representation. The 3 guides targeting each gene are shown separately with the gene-level result sized by cell number. **G.** Comparison of proportions of guides between the CROP-seq vector and cells in Perturb-seq as an indicator of gene ‘essentiality’. Left: Volcano plot of gene enrichment in the vector compared to cells. Significance was calculated by the paired Generalized Linear Model (GLM) framework implemented in EdgeR (see Methods). Most significant results are genes which are depleted upon knockdown (ie cause a defect in cell viability). Right: Bars of raw data (%) for the top most significantly depleted (“essential”) or enriched (“genes”) genes.

**Figure S2.**
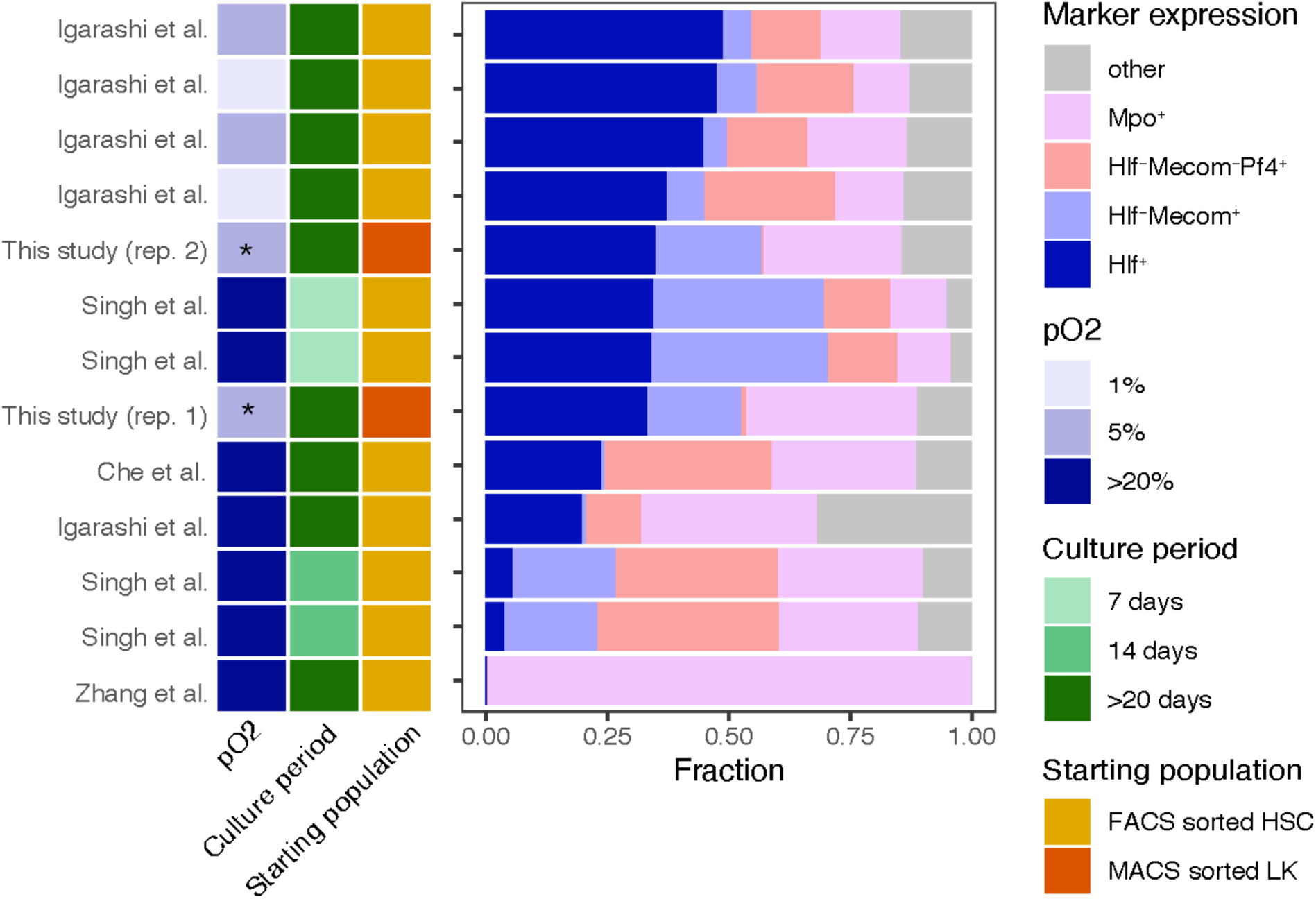
Comparison of cellular composition defined by transcriptomic marker genes between our study and others using HSPC expansion cultures. Related to Figure 1.

**Figure S3.**
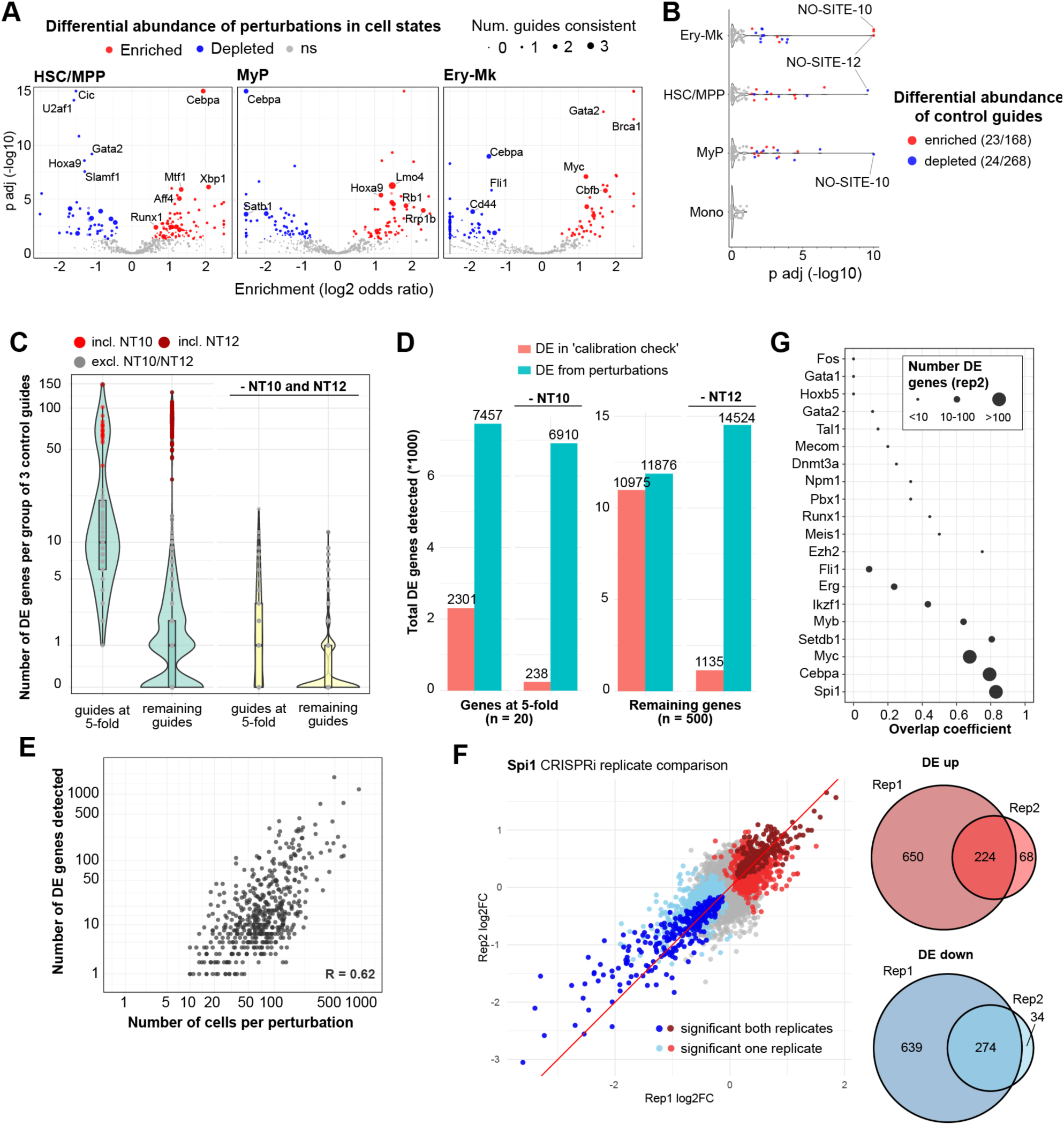
Perturb-seq differential abundance and differential expression (DE) analysis, related to Figure 1 and Supplementary Note. **A.** Volcano plots representing differential abundance of perturbations across three broadly defined cell states in the Perturb-seq data. Dot size represents the number of individual guide RNAs targeting that gene that were significant when tested separately. **B.** Differentially abundant non-targeting control guide RNAs in cell states, as calculated by Fisher’s exact test comparing counts of specific control guides compared to counts of remaining controls and corrected for multiple-testing (Benjamini-Hochberg). **C.** Numbers of perturbation-response gene pairs called as significant from the “calibration check” of SCEPTRE analysis (see Supplementary Note). Each point in the violins represents a test for differential expression between a random group of 3 non-targeting control guides and remaining controls. Analysis was run separately with guides included at 5-fold coverage and the remaining data set. Upon removal of NO-SITE-10 and NO-SITE-12 from the set of non-targeting control guides, many fewer pairs were called as significant in the calibration check. For C-G, hits are reported as significant after calculation of p-values by SCEPTRE’s permutation-based framework for differential expression testing and subsequent multiple testing correction (Benjamini-Hochberg). **D.** Total numbers of perturbation-response gene pairs called as significant in the ‘calibration check’ compared to ‘discovery analysis’ using the real perturbations in the data. As the numbers of DE tests are equal, this comparison acts as an estimate for the rate of false positives due to miscalibration of DE analysis. **E.** Scatter plot comparing the number of genes detected as differentially expressed downstream of a perturbation with the number of cells containing guides targeting the gene. Data is from replicate 1. **F.** Comparison of differential expression from *Spi1* knockdown between replicate 1 and replicate 2, in which fewer cells were profiled. DE data from both replicates is shared on Figshare. **G.** Quantification of the fraction of differentially expressed response genes in Replicate 2 that were also identified in Replicate 1 (overlap coefficient) for the 20 priority gene targets included at five-fold representation in the CRISPRi library.

**Figure S4.**
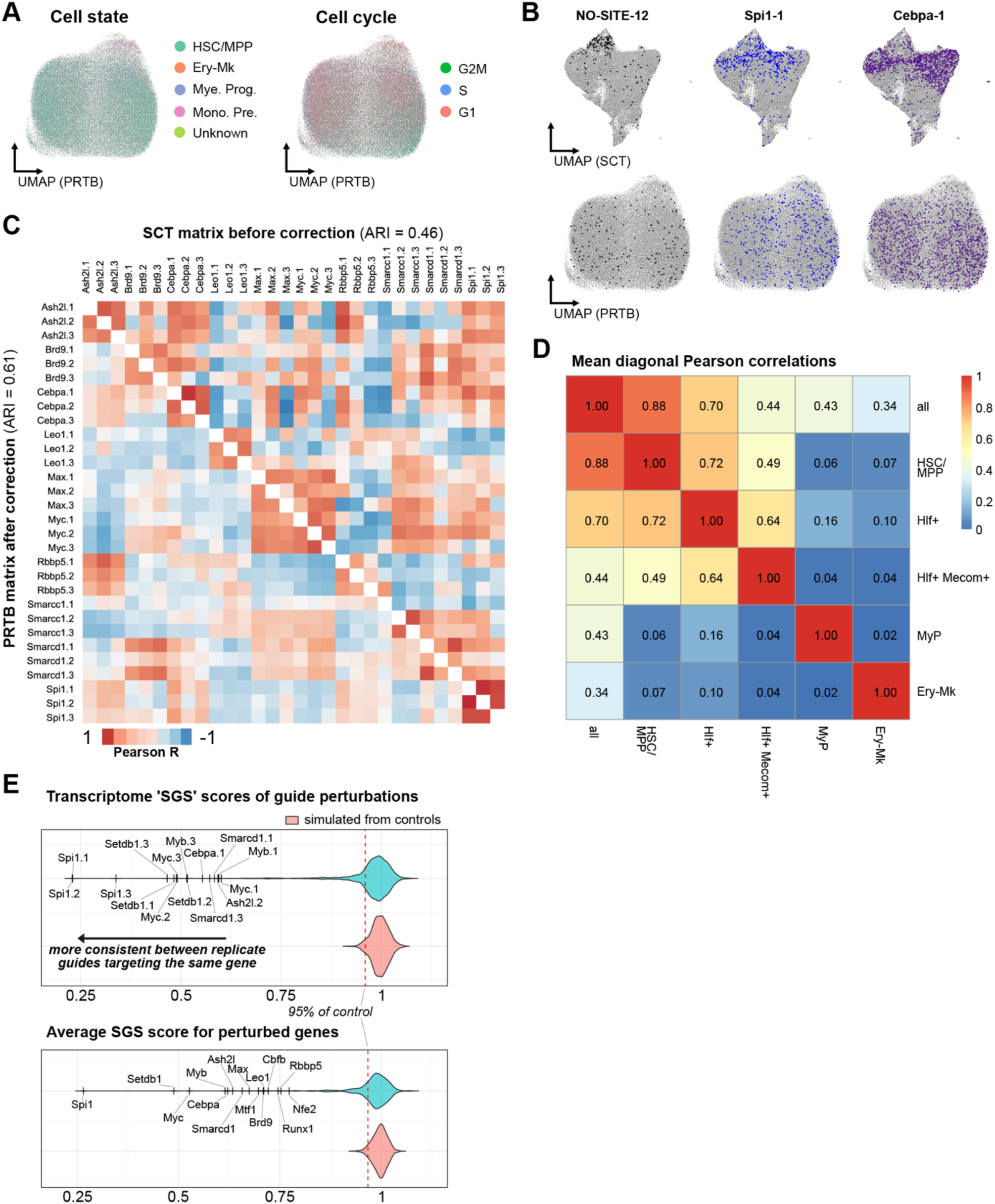
Neighborhood transformation masks clonal heterogeneity to aid discovery of consistently co-regulating perturbations, related to Figure 1. **A.** UMAP embeddings of single cells after neighborhood transformation, labelled by cell state and cell cycle phase annotations. **B.** Comparison of distribution patterns of cells containing example guide RNAs over single-cell UMAPs before (above) and after (below) neighborhood transformation. **C.** Heatmap of pairwise correlations calculated from “pseudobulks” of selected gene perturbations. Correlations without neighborhood transformation are plotted **above the diagonal**, and from neighborhood-transformed perturbation signatures **below the diagonal**. The Adjusted Rand Index (ARI) quantifies how well guides targeting the same gene are grouped in the same heirarchical clusters, with the number of clusters fixed as the number of genes. The color scale (Pearson R) is slightly exaggerated for visual effect. **D.** For each gRNA, pseudobulk PRTB scores were computed in subsets of the data made from different cell states, and correlations between RTPB scores per gRNA between cell types were computed. Heatmap shows the average (across gRNAs) correlation between pairs of cell states. **E.** Violin plots of Same-Gene-Similarity (SGS) scores calculated from comparisons of perturbation pseudobulks (see Methods, *Filtering for consistent perturbations*). Above: SGS scores of individual guide perturbations. Below: Average SGS scores over 3 guides per gene. The red vertical dashed line indicates a cut-off for “consistent” perturbations, defined as an SGS score lower than the 5th percentile of simulations of randomly grouped triplets of control guides.

**Figure S5.**
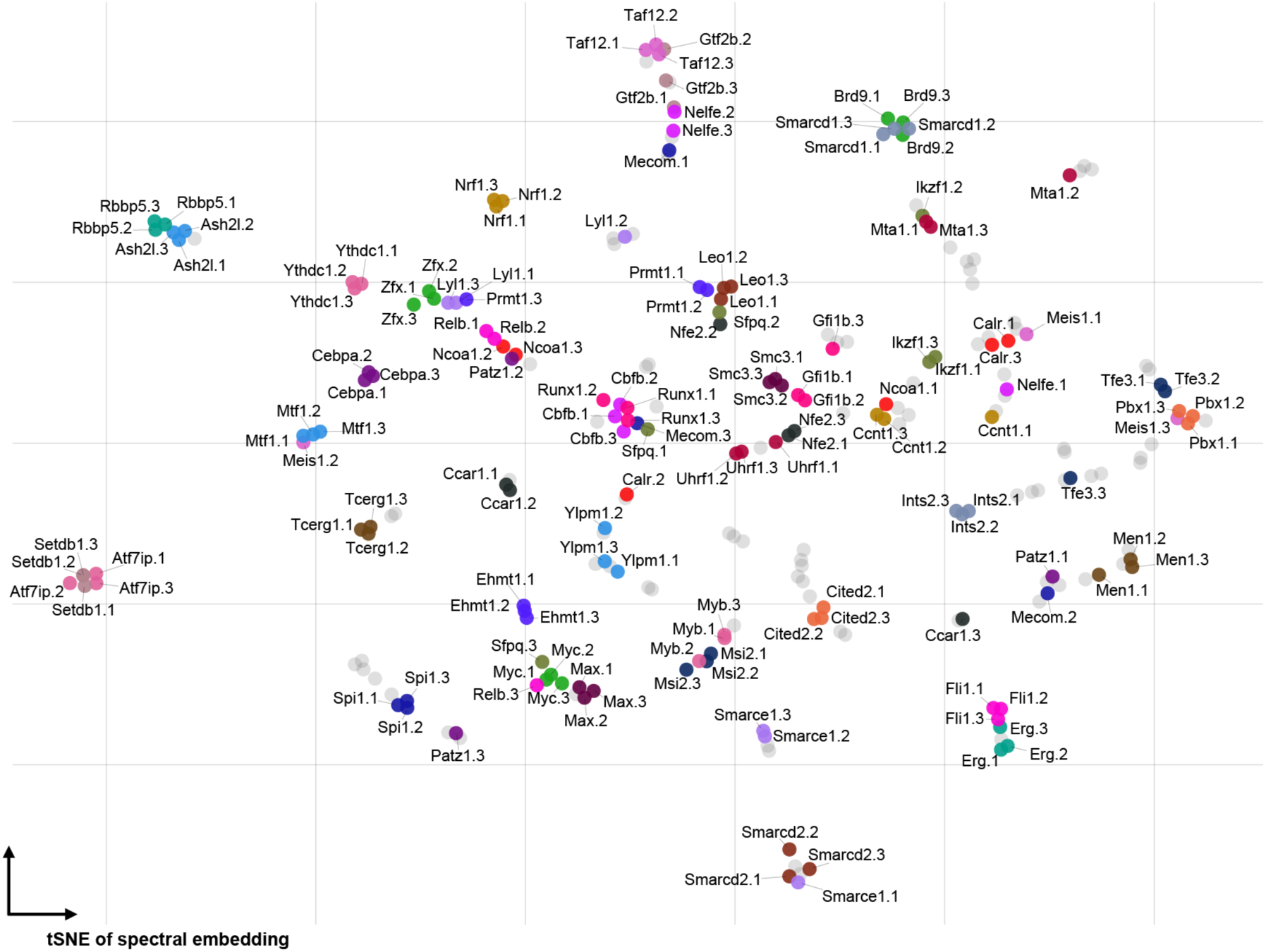
tSNE embedding of perturbations by similarity of effects on the HSPC transcriptome (co-regulat*ing* genes), labeling all perturbations with 3 guides per gene after consistency filtering. Related to Figure 1.

**Figure S6.**
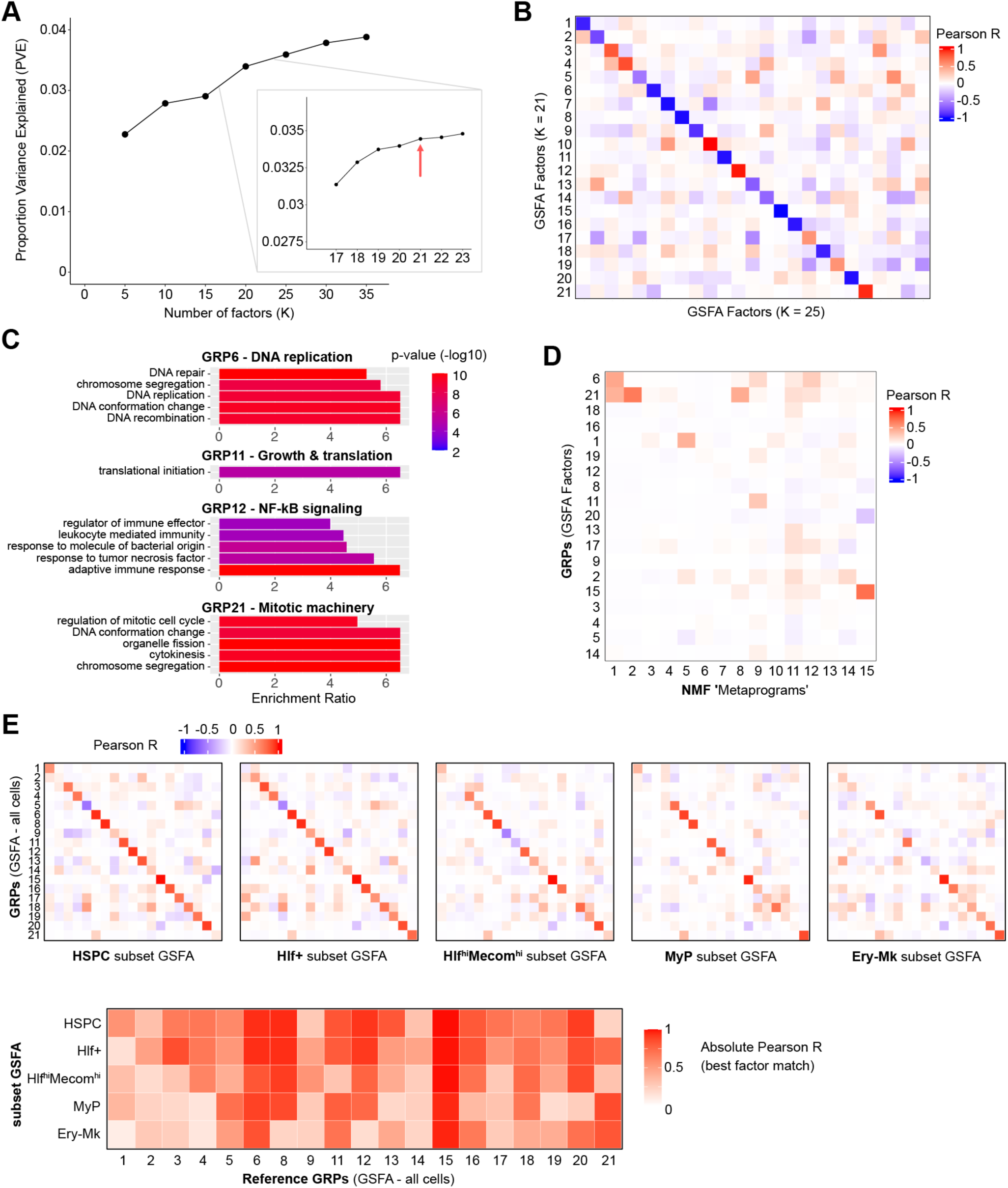
Guided Sparse Factor Analysis (GSFA) defines gene regulatory programs and their regulation, related to Figure 2. **A.** Proportion Variance Explained (PVE) by latent factors from GSFA using different values of the K parameter (the number of factors to infer). PVE was calculated as the fraction of variance explained by linear combination of factors over the total variance in the neighborhood-transformed single-cell gene expression matrix used as input for GSFA (see Methods). **B.** Heatmap of pairwise correlations between factors derived from GFSA with K=21 compared to K=25 (Pearson correlations across all gene weights in W matrices). Assignment of numbers to factors is arbitrary so are only labelled for the result we continued with for downstream GRP annotation and further analysis. **C.** Enrichment of GO terms from the “geneontology_Molecular_Function_noRedundant” database in lists of genes positively loaded to selected factors (GRPs), calculated by WebGestaltR Over-Representation Analysis (ORA). **D.** Heatmap of pairwise correlations between factors derived from GSFA and metaprograms of gene co-expression derived from NMF. Comparisons are between sparsified matrices with the same features set (genes). **E.** (above) Heatmap comparisons of factors inferred by GSFA across different cell states compared to factors inferred from the whole data (=’GRP’s). Factor labels are arbitrary so the x axes are reordered so highest correlations with GRPs lie along the diagonal. (below) Summary of best matches between GRPs and cell state factors (diagonals from above).

**Figure S7.**
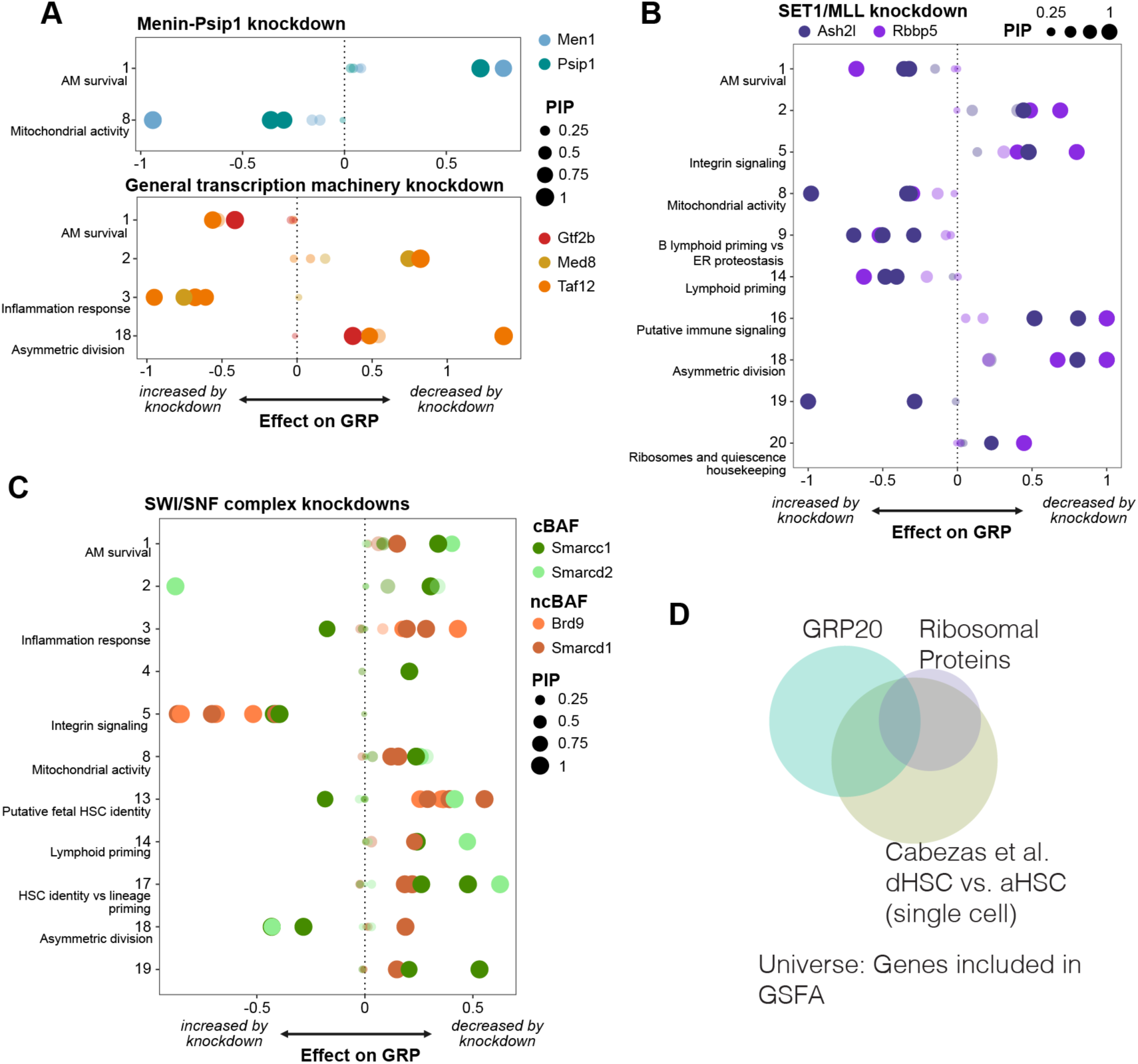
Further analysis of GRPs and their regulation, related to figures 2 and 3. **A-C.** Dot plots of the effects of various perturbations to GRPs. Colors of dots represent strength of effect and sizes represent the likelihood that the perturbation affects the GRP (Posterior Inclusion Probability). Only GRPs with 2 or more guide perturbations carrying PIP > 0.8 are shown in the plots. **D.** Venn diagram illustrating the overlap of GRP20, a dormant HSC signature^49^ and ribosomal protein genes

**Figure S8.**
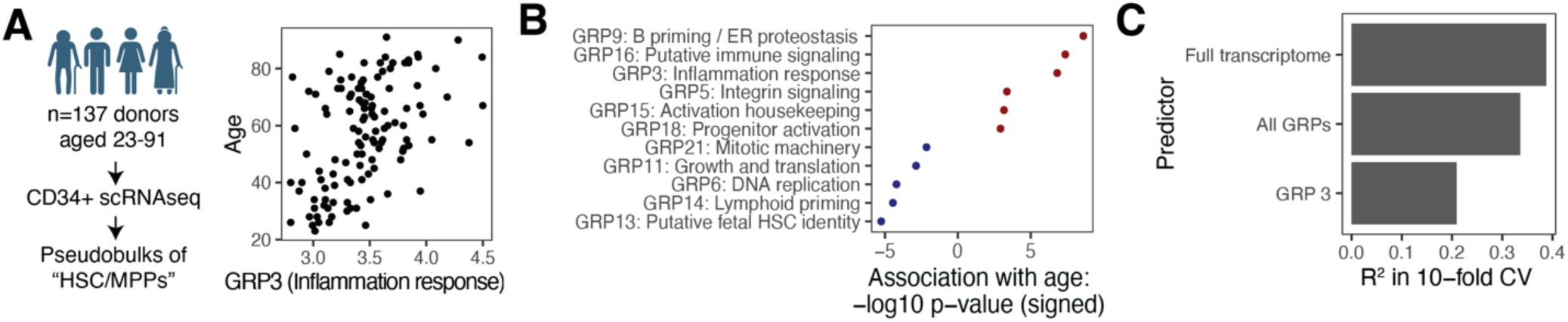
GRPs in human ageing, related to Figures 3 and 4. **A.** Decomposition of HSC transcriptomes from an ageing cohort^52^ demonstrates increase of GRP3 in age. **B.** Association of different GRPs with human age. Only significant (FDR<0.1) associations are shown. See Methods, section *Decomposition of HSC transcriptome data into GRPs* for detail. **C.** Bar chart quantifying fraction of age variance explained by different predictors.

**Figure S9.**
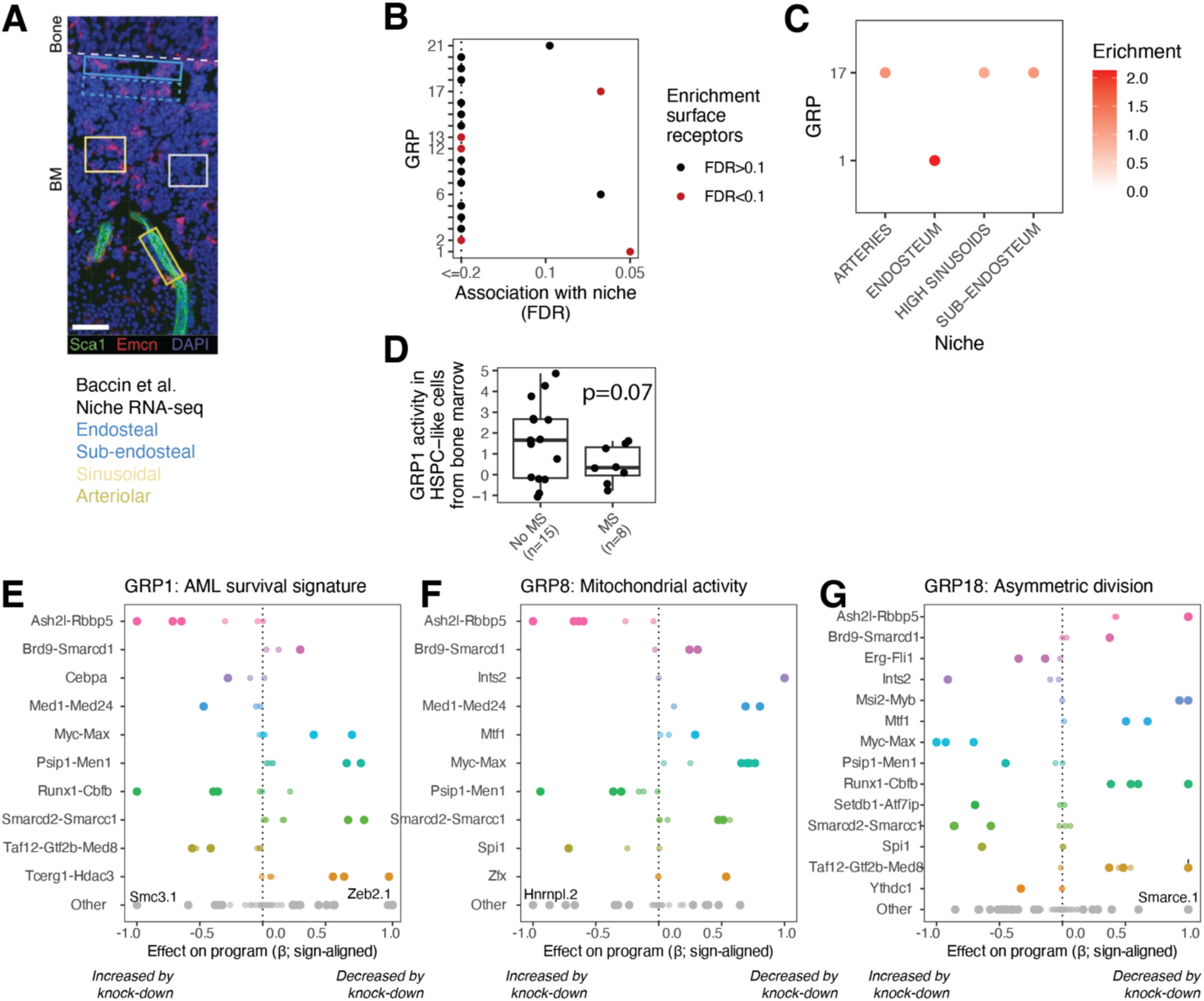
GRPs in acute myeloid leukemia. Related to Figure 4. **A.** Image illustrating the spatial transcriptomic dataset used for panel B and C, adapted from^87^. Bone regions were selected by Laser Capture Microdissection, and resulting transcriptome data was decomposed into GRPs. **B.** Plot depicting the association of GRPs with niche, and the enrichment of surface receptor genes on GRPs. P-values for niche association (x axis) were computed using Anova. P values for enrichment of surface receptor genes (color coded) was computed using a hypergeometrical test. **C.** Enrichment of GRPs in different niches. Enrichment (color coded) stems from a linear model of inferred GRP activity as a function of niche. All associations shown are significant (FDR < 10%). **D.** Association of GRP1 with extramedullary disease. HSPC-like cells were selected from a single-cell study of bone marrow from AML patients with and without myeloid sarcoma^59^. GRP activity was determined by decomposition. **E-G.** Dot plot illustrating effects of knockdown on GRP1 (E), GRP8 (F) and GRP18 (G).

## Supplementary Table Legends

**Table S1.** Selection of genes to target for CRISPRi Perturb-seq in HSCs. Tab 1 of the spreadsheet collects information for each gene in the mouse genome and assigns points according to the ranking criteria (see Methods: *Selection of genes in CRISPRi Perturb-seq)*. Tab 2 contains the final list of 520 genes included in the experiment. Tab 3 lists the sequences of the guide RNAs and oligos ordered from Twist for pooled cloning of the library (see Methods: *Guide RNA library design and cloning into CROP-seq-Puro-eGFP*).

**Table S2.** HDBSCAN clustering of tSNE embedding of co-regulating perturbations, related to Figure 1G and 2G. Numerical names of clusters (Figure 2G x-axis) are in the “cluster_merge” column.

**Table S3.** Lists of genes positively and negatively loaded to each GRP. Genes were filtered by posterior inclusion probability > 0.8 and are listed in descending order by weight of loading.

## Supplementary Note: Differential expression (DE) analysis, related to Figure S3

### Evaluation of calibration of differential expression with SCEPTRE

We chose to use SCEPTRE^33^ for differential expression analysis. A key feature of this R package is a “calibration check” designed to assess how well calibrated DE association testing is for a given Perturb-seq data set. We reasoned that applying this to our data could help estimate the extent of potential false positives arising from heterogeneous patterns of guide RNAs driven by clonal proliferation rather than perturbation effects. The calibration check is implemented by sampling random groups of non-targeting control guides (matched to the number of guides per gene in the experimental design, in our case 3) and tests for differential expression between this group and the remaining controls. It is explained in more detail in Barry et al, *Figure 1*, and here: https://timothy-barry.github.io/sceptre-book/run-calibration-check.html.

To apply SCEPTRE, we first ensured that random groups of 3 controls would be equivalent to gene-level perturbations by separating our data into the set of guide RNAs included at 5-fold (3 x 20 targeting guides + 20 non-targeting controls) and the set of “remaining” guides. We set sparsity thresholds to restrict analysis to response genes detected in sufficient numbers of cells, and then performed association testing for differential expression. SCEPTRE’s framework uses permutations of a negative binomial generalized linear model (GLM) to account for cell-specific confounding factors such as library size or batch/biological replicate, and also corrects p-values for multiple testing (more detail is provided in Barry et al^33^, *Methods*). Calibration of DE with respect to potential false positives is estimated by performing an equal number of “discovery tests” (tests for differential expression between a gene perturbations and response genes) and “calibration tests” (using the aforementioned negative control sets).

When applied to our data, we initially noted high numbers of significant associations from the calibration check for both 5-fold and remaining control guides (Figure S3C, left violins). Further inspection revealed that calibration groups containing NO-SITE-10 and NO-SITE-12, the two specific control guide RNAs most differential enriched between cell states (see Figure S3B), were responsible for the vast majority of significant associations in the calibration check (Figure S3C,D). Removal of these two guide RNAs from the control sets used for DE testing drastically reduced the numbers of significant associations detected per control group (Figure S3C, right violins) and resulted in a low overall estimated rate of false positives (Figure S3D; 0.034 for 5-fold genes and 0.078 for remaining genes), demonstrating good calibration for DE testing over the rest of the data.

### Estimation of how many gene perturbations are at risk of inflated false positives

Importantly, removing NO-SITE-10 and NO-SITE-12 does not address the underlying issue that many targeting guide RNAs also exhibit cell state biases (whether arising from perturbation or from clonal heterogeneity) and therefore remain susceptible to inflated false positives in DE. However, the observation from the calibration check that only 2 out of 120 controls are strongly affected suggests that false positives from clonal effects likely only impacts a minority of gene perturbations (i.e. DE results for most gene perturbations are reliable). We can estimate the fraction of genes affected as follows:

1. If *p* is the underlying probability a guide produces inflated false positives, and *K* is the number of affected guides out of *n* = 120 controls, then:

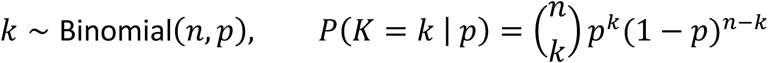

With *k* = 2, the empirical estimate is:

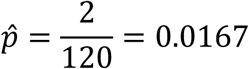

Because k is small, this estimate of *p* carries large uncertainty. The 95% Clopper-Pearson confidence interval calculated by inverting the binomial is:

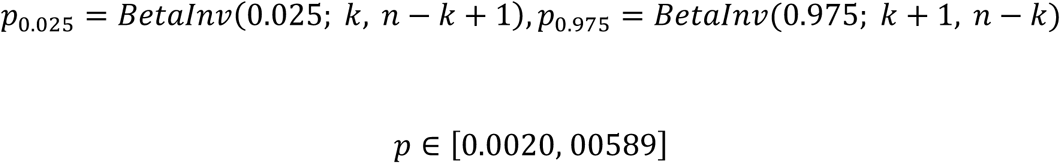
2. If we assume that any of the 3 guides per gene produces inflated false positives for a gene-level calculation, then the probability for genes (*q*) is given by:

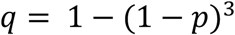

This produces estimates of:

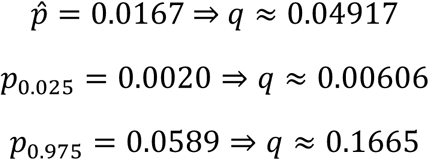
3. Multiplying *q* by the 520 genes in the data set:

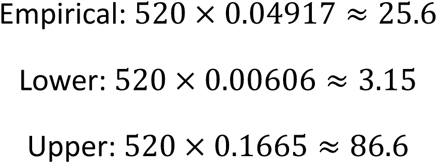

Thus, ∼26 genes are expected to affected by inflated false-positive DE, but this estimate has high uncertainty (CI_95%_ from ∼3 to ∼87).

### Consistency between 3 guides targeting each gene

As an additional indicator of which DE results to trust, we performed SCEPTRE on a guide-by-guide basis for the 3 guides targeting each gene. The logic behind this is that consistent effects across all three guide ‘replicates’ provides stronger evidence that a response gene is differentially expressed upon perturbation. Consistency was quantified by three metrics:

1. Sign agreement (*A*), the fraction of the 3 guides that match the direction of the overall fold-change of the gene perturbation.
2. A composite “consistency score”, defined as:

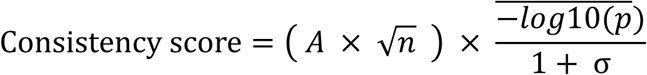

where

- *A* is the sign agreement (above)
- *n* is the number of guides targeting the guide (typically 3, but can be 1 or 2 if insufficient cells for some guides)
- 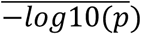 is the mean of the p values (-log10) across guides
- σ is the standard deviation of fold-changes (log2) across guides
3. A meta-analysis p value (p_meta_adj). This was generated by first calculating signed Z-scores from the p-values per guide (*p_i_*):

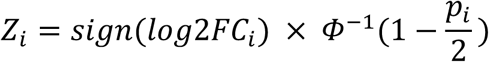

Where *Φ*^−1^ denotes the inverse cumulative distribution function of the standard normal distribution.

Signed Z-scores were combined using Stouffer’s method and converted back to p-values and corrected for multiple testing (Benjamini-Hochberg).

We provide the per-guide results and these scores as additional columns in the **full table of differential expression results** uploaded to Figshare.

### Overlap of DE results in a biological replicate experiment

Another approach to increase confidence in differential expression results is biological replication. We performed a biological replicate of the CRISPRi Perturb-seq screen, using HSPC cultures established from different mice and a distinct lentivirus preparation, but otherwise following the same experimental protocol as Replicate 1. A smaller number of cells (19,702) were profiled with an Evercode Mini kit (Parse Biosciences) for scRNA-seq by combinatorial barcoding.

The capacity to detect differential expression in Perturb-seq is strongly associated with the number of cells per perturbation (Figure S3E). Thus, replicate 2 was underpowered for DE detection for the majority of the 520 perturbations. However, overlap between replicates was strong for perturbations with sufficient cell coverage, with most DE hits in Replicate 2 also detected in Replicate 1. We demonstrate this for the example of knockdown of *Spi1* in Figure S3F and across priority gene targets included at five-fold representation in the CRISPRi library (Figure S3G, overall overlap coefficient of 0.716),

An annotated Seurat object containing all cells of Replicate 2 and the SCEPTRE differential expression results for 5-fold covered genes in Replicate 2 are shared on Figshare.

